# ATAC-seq reveals megabase-scale domains of a bacterial nucleoid

**DOI:** 10.1101/2021.01.09.426053

**Authors:** Michael D. Melfi, Keren Lasker, Xiaofeng Zhou, Lucy Shapiro

## Abstract

Here we adapted ATAC-seq to probe chromosome accessibility of bacterial cells. We found that the chromosome of Caulobacter crescentus is composed of eight differentially compacted regions we name Chromosomal Accessibility Domains (CADs). This domain structure is depended on the cell cycle stage, DNA gyrase activity, and the nucleoid-associated protein (NAP) GapR, but not on the function of SMC. We show the chromosome is punctuated by four <u>h</u>ighly transposase-<u>in</u>accessible transcribed regions (HINTs). The HINTs include Caulobacter’s ribosomal RNA clusters and its largest ribosomal protein gene cluster. Further, we show that HINTs are also formed by rDNA in E. coli and provide evidence that their high levels of transcription do not strictly govern their formation. Overall, this work argues that physical forces, including those created by the activities of DNA gyrase and specific NAPs, significantly contribute to bacterial nucleoid structure at the megabase scale.

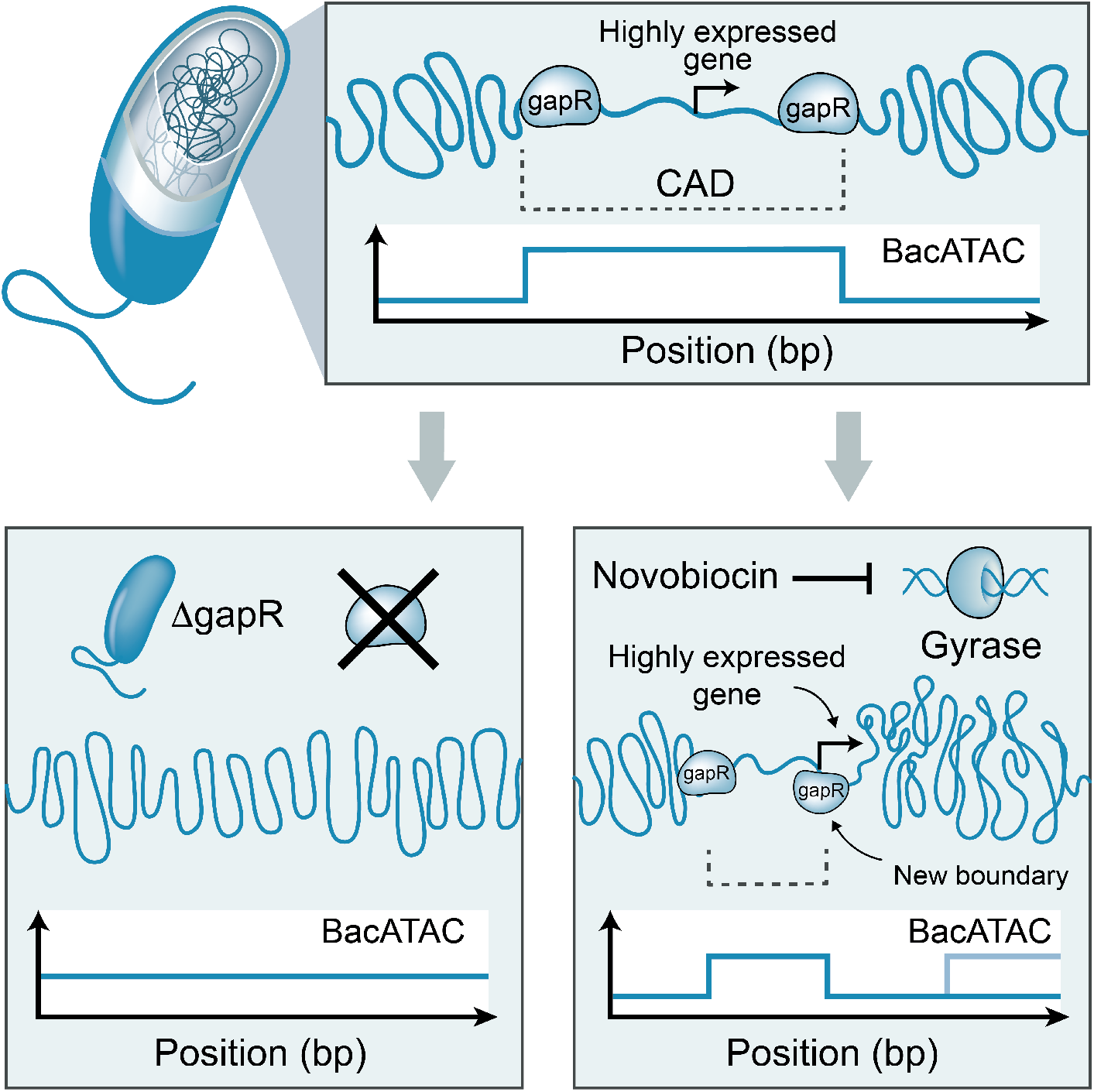

**Significance:** In bacteria, chromosomal DNA is highly compacted and organized. Many forces contribute to bacterial DNA compaction, including the transcription, DNA replication, and the activities of topoisomerases and nucleoid-associated proteins. At the megabase scale, the resulting chromosome structure is important for coordinating cell cycle events; for example, in *E. coli* the improper structuring of a Mb-scale nucleoid domain leads to errors in the fidelity of chromosome segregation. It was previously unknown whether Mb-scale regions of a bacterial chromosome could be differentially compacted, and which factors might contribute to this spatial variation in compaction. Our work provides a novel method for measuring global chromosome compaction in bacteria. By applying this method in the bacterium *Caulobacter crescentus*, we show that the *Caulobacter* nucleoid’s compaction varies between megabase-scale domains. We also show that DNA gyrase and the nucleoid-associated protein GapR are key players in shaping the uneven compaction of the chromosome.

## Introduction

Despite lacking nucleosomes, the bacterial nucleoid is highly organized (1). At the ∼10kb scale, bacterial DNA is stochastically looped into transient topological domains – DNA regions in which local DNA supercoils are temporarily trapped (2). High-resolution contact maps revealed that the bacterial chromosome, of Escherichia coli (3), Caulobacter crescentus (4), and Bacillus subtilis (5) are segmented into ∼100kb highly self-interacting regions called chromosomal interaction domains (CIDs). In E. coli, these CID coalesce into four Mb-scale macrodomains and two non-structured regions (3, 6). These macrodomains are distinct regions that display uniform physical behaviors including diffusivity and interaction frequencies (7). Here, we have asked whether there are Mb-scale domains in the Caulobacter nucleoid.

ATAC-seq (<u>A</u>ssay for <u>T</u>ransposase-<u>A</u>ccessible <u>C</u>hromatin using <u>seq</u>uencing) assesses nucleosome occupancy on DNA in eukaryotic cells by probing accessibility of DNA to transposase enzymes in intact nuclei (8). We developed Bac-ATAC (Bacterial ATAC-seq), an adaptation of ATAC-for bacterial cells. Using Bac-ATAC we interrogated the accessibility of the *Caulobacter* chromosome at single base-pair resolution. We found that the *Caulobacter* chromosome is assembled into eight Mb-scale regions we name CADs (<u>C</u>hromosomal <u>A</u>ccessibility<u>D</u>omains). These domains are characterized by unique average accessibilities. We show that the average accessibility of a CAD encodes information about the relative amount of cellular space it occupies and therefore its physical level of compaction. Therefore, we argue that CADs are equivalent to regions of unique average compaction, and we propose that they behave as Mb-scale topological domains. DNA gyrase and the nucleoid-associated protein (NAP) GapR play non-redundant roles in maintaining wild type differential compaction of CADs. Finally, we revealed four Highly Transposase-Inaccessible Transcribed (HINT) regions of the chromosome, three of which are ribosome-associated. We show that in *E. coli*, HINTs are formed by DNA regions that encode ribosomal genes. One of the four *Caulobacter* HINTs became accessible to transposon insertion upon infection by phage, despite no detectable change in HINT expression, implying that HINT formation is not controlled strictly by transcription extent. Overall, our assessment of megabase-scale chromosome compaction in *Caulobacter* leads to a restratification of the forces known to modulate bacterial nucleoid structure. While previous work has shown that transcriptional forces contribute to chromosome structure on short length scales (*i*.*e*., *via* CID formation), on Mb-length scales non-transcriptional forces including those generated by the activities of NAPs and DNA gyrase emerge as key shapers of bacterial nucleoid structure.

## Results and Discussion

### ATAC-seq for bacterial cells

We developed Bac-ATAC, a bacteria-compatible ATAC-seq protocol (**Material and Methods 1, SI Methods 1**), to measure variations in transposase-accessibility of the *Caulobacter* chromosome in intact cells. The protocol includes fixation, permeabilization, transposition, reverse crosslinking, and library preparation steps. Performance of Bac-ATAC on milliliters of midlog phase *Caulobacter* cultures yielded genome-wide transposon insertion frequency data in high-throughput and at single bp-resolution from a single experiment. The insertion data was then normalized for sequence bias and copy number by comparing insertion frequencies in a given sample to those generated by performing the same method on purified genomic DNA (**Materials and Methods 2**).

In a mixed population (**Figure 1a**) of *Caulobacter* cells, we mapped more than 27 million insertions, which covered over 80% of its circular 4 Mb chromosome (**SI Table 4**). The chromosome appeared largely accessible to transposase *in vivo*, with the longest continuous DNA stretch avoiding insertion stretching 33bp. This is in agreement with findings in *E. coli*, in which the chromosome is generally accessible to restriction enzymes (2).s

**Figure 1.**
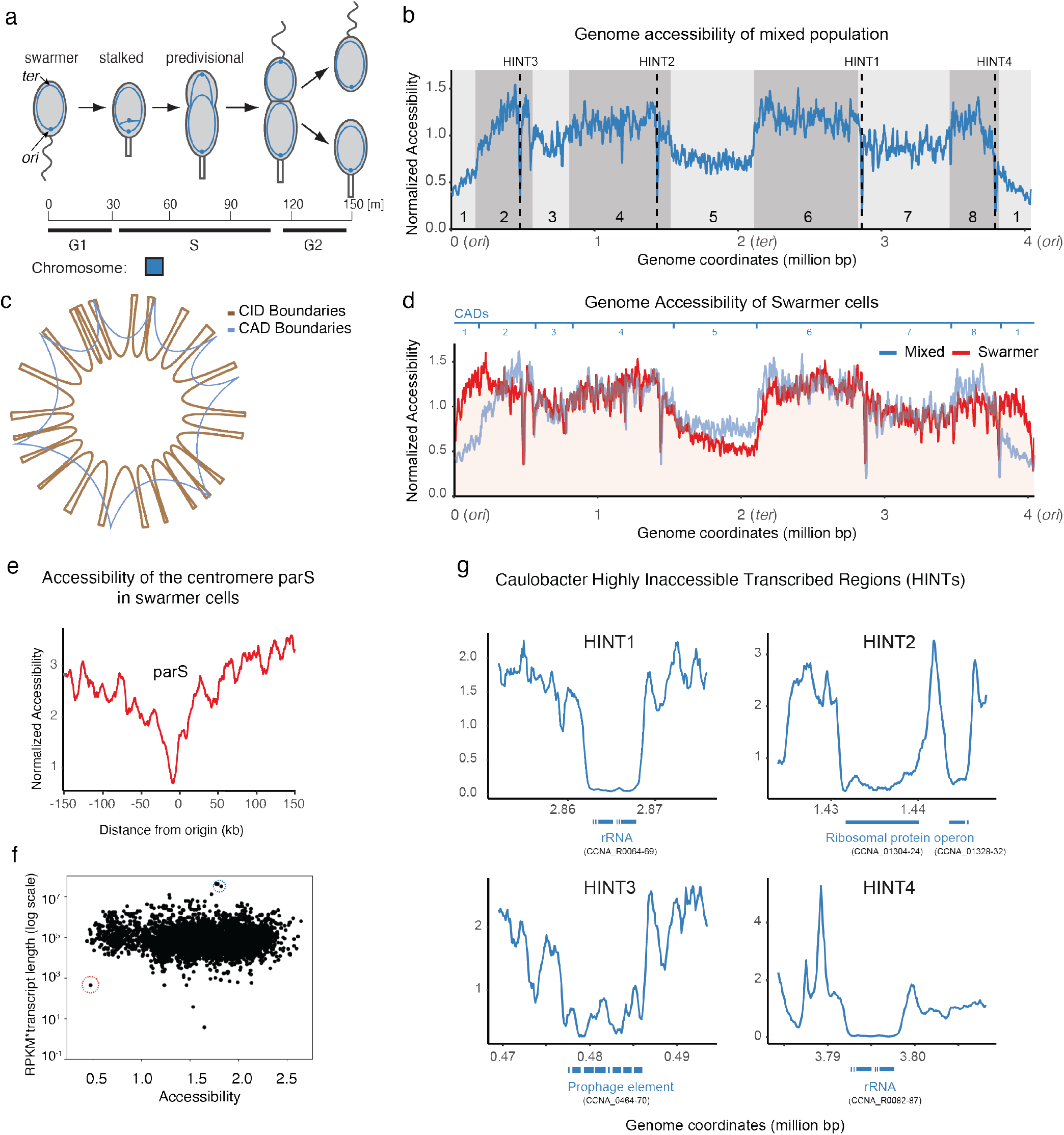
Caulobacter chromosome is divided to accessibility regions. **a**. Schematic of the dimorphic cell cycle of *Caulobacter crescentus*. The origin and terminus of the chromosome are labeled as “ori” and “ter” respectively. During the cell cycle, the motile, non-replicating swarmer cell first differentiates into a replication-competent stalked cell before dividing asymmetrically into two morphologically distinct cell types. **b**. Linear plot of chromosome accessibility determined from Bac-ATAC data mixed population of *Caulobacter* cells (blue). The X-axis starts at the origin (left), proceeding clockwise around the chromosome, through the terminus (middle), and returning to the origin (right). Black dotted lines represent global minima of accessibility. CADs are numbered 1-8, and even-numbered Chromosomal Accessibility Domains (CADs) determined using mixed population accessibility data are shaded grey. **c**. CID boundaries (brown) and CAD boundaries (blue) overlaid in polar coordinates. **d**. Overlay of mixed population (blue) and swarmer (red) ATAC-seq data in Cartesian coordinates. Axes and CAD numbering are the same as in (b). **e**. Bac-ATAC data, focused on the *parS* centromeric region of the *Caulobacter* swarmer chromosome. The origin is at x = 0. *parS* is an inaccessible region of the swarmer cell nucleoid (**SI Note 12**). **f**. Log-Log plot of Expression vs. accessibility for all genes in Caulobacter. The most inaccessible gene is a HINT-proximal ferredoxin protein CCNA_03639 (red circle). The most highly expressed mRNA in Caulobacter, *rsaA*, has average accessibility (blue circle). **g**. WT ATAC accessibility data for HINT regions. Blue boxes represent Highly Inaccessible Transcribed Region genes. Each box represents individual transcriptional units identified in (15) for HINTs 2,3 and individual genes for HINTs 1,4. Specific gene names are in **Table 1**. Data shown was smoothened to 1kb. Blue boxes are not precisely to scale with respect to one another. Data is normalized to gDNA control data; smoothening was done using sliding window averaging with a window size of 1kb (**SI Methods 11**).

### The Caulobacter chromosome contains Mb-scale regions that have characteristic accessibility levels

To find trends of insertions at longer length scales we smoothened the Bac-ATAC insertions data across 10 kb windows and searched for points of sudden discrete changes in the data. This analysis revealed that the *Caulobacter* chromosome is structured into eight megabase-scale (>100kb) regions that feature differential, characteristic average levels of accessibility (**Figure 1b**). We name these regions Chromosomal Accessibility Domains (CADs) and refer to the change-points that delineate these regions as CAD boundaries. We note that while all CADs contain two or more of *Caulobacter’*s previously discovered Chromosomal Interacting Domains (CIDs), CAD boundaries do not precisely overlap CID boundaries; only three of the eight CAD boundaries are found within CID boundaries (**Figure 1c**; **SI Figure 1**).

**Table 1:** Inaccessible Transcripts

| Transcript | Transcript Annotation | Rank** |
| --- | --- | --- |
| **HINT 1: rRNA (7kb, Extrema coordinate = 2863911, 7kb)** |  |  |
| CCNA_R0064 | tRNA fMet |  |
| CCNA_R0065 | 5S ribosomal RNA |  |
| CCNA_R0066 | 23S ribosomal RNA |  |
| CCNA_R0067 | tRNA-Ala |  |
| CCNA_R0068 | tRNA-Ile |  |
| CCNA_R0069 | 16S ribosomal RNA |  |
| **HINT 2: rRNA (7kb, Extrema coordinate = 1435826, 9.5kb and 2.5kb)** |  |  |
| CCNA_01304 | hypothetical protein | 2 |
| CCNA_01305 | SSU ribosomal protein S10P | 7 |
| CCNA_01306 | LSU ribosomal protein L3P | 32 |
| CCNA_01307 | LSU ribosomal protein L1E-L4P | 30 |
| CCNA_01308 | LSU ribosomal protein L23P | 38 |
| CCNA_01309 | LSU ribosomal protein L2P | 31 |
| CCNA_01310 | SSU ribosomal protein S19P | 33 |
| CCNA_01311 | LSU ribosomal protein L22P | 49 |
| CCNA_01312 | SSU ribosomal protein S3P | 35 |
| CCNA_01313 | LSU ribosomal protein L16P | 34 |
| CCNA_01314 | LSU ribosomal protein L29P | 22 |
| CCNA_01315 | SSU ribosomal protein S17P | 42 |
| CCNA_01316 | LSU ribosomal protein L14P | 11 |
| CCNA_01317 | LSU ribosomal protein L24P | 14 |
| CCNA_01318 | LSU ribosomal protein L5P | 3 |
| CCNA_01319 | SSU ribosomal protein S14P | 18 |
| CCNA_01320 | SSU ribosomal protein S8P | 28 |
| CCNA_01321 | LSU ribosomal protein L6P | 23 |
| CCNA_01322 | LSU ribosomal protein L18P | 17 |
| CCNA_01323 | SSU ribosomal protein S5P | 45 |
| CCNA_01324 | LSU ribosomal protein L30P | 66 |
| CCNA_01328 | SSU ribosomal protein S13P | 25 |
| CCNA_01329 | SSU ribosomal protein S11P | 8 |
| CCNA_01330 | DNA-directed RNA polymerase alpha chain | 53 |
| CCNA_01332 | LSU ribosomal protein L17P | 64 |
| **HINT 3: prophage element (7kb, Extrema coordinate = 481472, 9.5 kb)** |  |  |
| CCNA_0464 | Undecaprenyl-phosphate sugar phosphotransferase-related protein | 563 |
| CCNA_0465 | UDP-galactopyranose mutase | 146 |
| CCNA_0466 | glycosyltransferase | 196 |
| CCNA_0467 | oligosaccharide translocase/flippase | 272 |
| CCNA_0468 | UDP glucuronic acid epimerase-related protein | 682 |
| CCNA_0469 | GT1-family glycosyltransferase | 256 |
| CCNA_03922 | hypothetical protein | 392 |
| CCNA_00470 | O-antigen polymerase | 502 |
| **HINT 4: rRNA (7kb, Extrema coordinate = 3796257, 6 kb)** |  |  |
| CCNA_R0082 | tRNA fMet |  |
| CCNA_R0083 | 5S ribosomal RNA |  |
| CCNA_R0084 | 23S ribosomal RNA |  |
| CCNA_R0085 | tRNA-Ala |  |
| CCNA_R0086 | tRNA-Ile |  |
| CCNA_R0087 | 16S ribosomal RNA |  |

\* the ribosomal protein operon-associated HINT is actually separated into two inaccessible regions in close proximity.
\*\* Average mRNA expression rank: RPKM in PYE mixed population; only reported for protein-coding genes. The non-protein coding genes listed are assumed to be much more highly expressed than any mRNA.

### CAD structures depend on the Caulobacter cell cycle stage

During the *Caulobacter* cell cycle, each cell division is asymmetric, yielding a morphologically distinct swarmer cell and a stalked cell. Chromosome replication initiates in the stalked daughter cell, but the swarmer daughter cannot initiate replication until it differentiates into a replication competent stalked cell (**Figure 1a**). Bac-ATAC analysis of swarmer cell populations showed that CAD structure in this cell type is largely similar to that of the mixed population (**Figure 1d**). However, a notable exception is CAD 1, containing the centromeric locus *parS* as well as the origin of replication. This entire CAD is distinctly more accessible in the swarmer cell population. Despite this difference, the *parS* locus remains uniquely transposase-inaccessible (i.e., sustaining few Bac-ATAC insertions) in the swarmer (**Figure 1e**).

### Genome accessibility does not correlate with transcriptional activity

We predicted that very highly expressed genes would appear particularly inaccessible to transposase, in part due to potential blockage by RNA polymerase machinery. To test this prediction, we measured the accessibility of DNA segments encoding *Caulobacter* transcripts. **Figure 1f** shows that transcript activity and degree of accessibility are not correlated (R^2^ = 0.070, **SI Figure 2**). Even very highly transcribed regions of the chromosome did not appear particularly inaccessible to transposase; for example, the *rsaA* locus, which encodes the most highly abundant mRNA in *Caulobacter* in rich media, displayed an average level of accessibility to transposase (**Figure 1f**). We therefore concluded that a genomic locus’s accessibility to transposase is not reflective of its transcriptional extent. However, we also uncovered a small number of notable exceptions, which we describe next.

### The nucleoids of both *E. coli* and *Caulobacter* contain Highly Transposase-Inaccessible transcribed regions, the majority of which are ribosome-associated

**Figure 1** shows that four exceptionally inaccessible loci visibly punctuate the *Caulobacter* chromosome (four black dotted lines in **Figure 1b**). Because these global minima of accessibility precisely overlap transcribed operons, we name these regions <u>H</u>ighly <u>In</u>accessible and <u>T</u>ranscribed regions (HINTs) (**Figure 1g**). The HINTs span from 2.5 to 9 kb in length. Three of the four HINTs are ribosome-associated in function; they include both of the *Caulobacter* ribosomal RNA operons and the largest ribosomal protein gene cluster (**Figure 1g, Table 1**). Accordingly, HINTs include some of the very longest and most highly expressed operons in *Caulobacter* (9).

**Table 2:** Locations of HINTs identified in *E. coli*

| Feature | Coordinate(s) (bp)* |  |  |  |  |  |  |
| --- | --- | --- | --- | --- | --- | --- | --- |
| *rrnH* | 229440 | 1134061 | 1400269 | 2015613 |  |  |  |
| *rrnG* | 2724912 | 2812722 | 3065848 | 3310627 |  |  |  |
| *rrnD* | 3422316 | 3547237 | 3624533 | 3717563 | 3809218 | 3881759 | 3921783 |
| *rrnC* | 3944412 |  |  |  |  |  |  |
| *rrnA* | 4037477 | 4080365 |  |  |  |  |  |
| *rrnB* | 4169138 |  |  |  |  |  |  |
| *rrnE* | 4210627 | 4468548 |  |  |  |  |  |

\*bp coordinate identified as a local minimum in smoothened Bac-ATAC data (See Materials and Methods for additional information). HINTs are comprised of these central points as well as the surrounding genomic areas.

**Table 3:** CAD Boundaries under novobiocin treatment

| novo CAD ID | CAD Boundary coordinate | Associated CID (if any) | Accessibility change<sup>°°</sup> | Direction of transcription | Genes located at novo-specific CAD boundary ** | Function | highly expressed in PYE ° | Highly expressed gene identified in ref. [4] | Do identified highly expressed genes match? | Is expression change novobiocin-induced *** | Fold change in expression | Long operon ‡ | Near WT CAD boundary †† | Change in boundary shape ‡‡ | In Fig. 2? |
| --- | --- | --- | --- | --- | --- | --- | --- | --- | --- | --- | --- | --- | --- | --- | --- |
| 1 | 51066 |  | increase | left | CCNA_000031 to CCNA_00060 |  |  |  |  |  |  | yes |  |  |  |
| 2 | 167461 | 2 | decrease | right | **gyrB (CCNA_00159)** | gyrase subunit B |  | CCNA_00161, CCNA_00162 | no † | yes | 8.5 |  | yes | inverts | yes |
| 3 | 391261 | 3* | increase | left | **CCNA_00374 operon (ATP synthase)** |  | yes | CCNA_00340, CCNA_00341 | no |  |  |  |  |  | yes |
| 4 | 447556 |  | decrease | right | CCNA_00439 to CCNA_00451 |  |  |  |  |  |  | yes |  |  |  |
| 5 | 472035 |  |  |  | artifact of HINT |  |  |  |  |  |  |  | yes |  |  |
| 6 | 487867 |  |  |  | artifact of HINT |  |  |  |  |  |  |  | yes |  |  |
| 7 | 522171 | 4 | decrease | right | **CCNA_00505, CCNA_00506** | cytochrome b, cytochrome c reductase | yes | L10P, L12P, RpoBB' | no |  | 505 = 1.5; 506 = 1.9; 507 = 1.8 |  |  |  | yes |
| 8 | 922346 |  | increase | left | CCNA_00850 (acrB2), CCNA_00849, CCNA_00851 | drug efflux system |  |  |  | yes | 849 = 6.9; 850 = 6.3; 851 = 7.6 |  |  |  |  |
| 9 | 949646 |  | decrease | right | CCNA_00878 | ATP-dependent RNA helicase |  |  |  | yes | 6.5 |  |  |  |  |
| 10 | 1086291 |  |  |  | no gene identified |  |  |  |  |  |  |  |  |  |  |
| 11 | 1124667 |  | decrease | right | CCNA_01034 | TonB-dependent outer membrane receptor |  |  |  | yes | 5.4 |  |  |  |  |
| 12 | 1157692 | 7 | decrease | right | **rsaA** |  | yes | rsaA | yes |  |  |  |  |  | yes |
| 13 | 1195549 |  |  |  | no gene identified |  |  |  |  |  |  |  | yes |  |  |
| 14 | 1237892 |  |  |  | CCNA_03890 to cckA | cckA |  |  |  |  | 4.4 |  |  |  |  |
| 15 | 1365706 |  |  |  | no gene identified |  |  |  |  |  |  |  | yes |  |  |
| 16 | 1427257 | 8 | decrease | right | **ribosomal protein cluster** |  | yes | ribosomal protein cluster | no |  |  | yes | yes |  |  |
| 17 | 1653905 |  |  |  | no gene identified |  |  |  |  |  |  |  |  |  |  |
| 18 | 1769833 |  | decrease | right | gyrA | gyrase subunit A |  |  |  | yes | 7.7 |  | yes | sharpens |  |
| 19 | 2174667 | 13 | increase | left | **nuo operon (CCNA_02012 to nuoA)** |  |  | S2P, trigger factor protein TIG | no |  |  | yes |  |  | yes |
| 20 | 2720585 | 16 |  |  | **CCNA_02565 to CCNA_02574** |  |  | S4P (CCNA_02596) |  |  |  |  |  |  |  |
| 21 | 2870256 | 17 | increase | left | **rRNA** |  | yes | rRNA | yes |  |  |  | yes | inverts |  |
| 22 | 2884698 |  | decrease | right | CCNA_02711 to CCNA_02721 |  |  |  |  |  |  |  |  |  |  |
| 23 | 2933619 |  |  |  | no gene identified |  |  |  |  |  |  |  |  |  |  |
| 24 | 3413217 |  |  |  | no gene identified |  |  |  |  |  |  |  |  |  |  |
| 25 | 3486085 | 20 | increase | left | **Ef-TU operon** |  | yes | Ef-TU operon | yes |  |  |  | yes | sharpens |  |
| 26 | 3535519 |  |  |  | no gene identified |  |  |  |  |  |  |  |  |  |  |
| 27 | 3776987 | 23 |  |  | artifact of HINT |  |  |  |  |  |  |  | yes | inverts |  |
| 28 | 3800718 | 23 | increase | left | **rRNA** |  | yes | rRNA | yes |  |  |  | yes | inverts |  |
| 29 | 3868151 |  | decrease | right | CCNA_03689 |  |  |  |  |  |  |  | yes | sharpens |  |
| 30 | 3988041 |  |  |  | CCNA_03806 to CCNA_03610 |  |  |  |  |  |  |  |  |  |  |

CID associate genes are **bolded and highlighted in blue**
°° Accessibility change moving from left to right along the chromosome
\* CID 3 boundary is actually ~1.5 kb away from change-point
\*\*Transcripts or genes located at novo-specific CAD boundary formation. Noteworthy genes found near accessibility change-points. "Artifact of HINT" means that this region appears as a change-point because of local change in accessibility at the edge of a highly inaccessible transcribed region; "no gene identified" means that there is no clear transcript or set of transcripts associated with this change-point. Blue shading implies the boundary is associated with a gene or set of genes that are highly expressed in novobiocin, with long operon, or with a HINT. No shading implies that the presence of the boundary cannot be explained by examining transcription of nearby genes alone.
‡ Here, long operons are defined as having twelve or more genes in a single operon, or having that many genes oriented in that direction in a row
† difference due to novobiocin-induced *gyrB* expression
°>=3000 rpkm in PYE [15]
\*\*\*at least one gene in operon has 2-fold or greater increase in gene expression and is also among the top 200 most highly expressed genes in novobiocin
‡‡ "Inverts" implies that the CAD boundary, or accessibility change-point, delineates the location of an increase in accessibility from left to right along the chromosome in WT conditions, but that during gyrase inhibition, it delineates a decrease in accessibility from left to right (or vice versa). "Sharpens" implies that the change-point is qualitatively more pronounced during gyrase inhibition than in WT conditions.

To test whether these specific ribosome-associated DNA regions in *E. coli* might also display inaccessibility, we performed Bac-ATAC on *E. coli* cells and measured the accessibility of ribosome-associated DNA. Strikingly, we see that each of the seven ribosomal RNA operons in *E. coli* (*rrnA-E,G,H*) rank among the twenty-one most highly inaccessible regions in *E. coli* (**SI Figure 3, Table 2**).

### Bac-ATAC measures chromosome compaction when analyzed at the megabase-scale

Many factors, including steric hindrance caused by protein binding, may affect bacterial chromosome accessibility on the 100-bp scale *in vivo* (10-12). Nevertheless, we predicted that on scales larger than 100kb, the accessibility of a chromosomal region would be inversely proportional to its level of compaction for geometric reasons (**SI Note 1**). To test this prediction, we performed simulated Bac-ATAC on high-resolution structural models of the *Caulobacter* chromosome (13) (**SI Methods 3**). These simulations demonstrated that accessibility and compaction are strongly negatively correlated, on both 15kb and 100kb scales. We therefore posit that Bac-ATAC, which directly measures chromosome accessibility, also provides information about the physical level of Mb-scale chromosome compaction.

### Inhibition of DNA gyrase yields new or altered CAD boundaries

DNA gyrase and topoisomerase IV (TopoIV), which add negative supercoils in an ATP-dependent manner, are the main drivers of DNA compaction in bacteria (14). To measure the effects of these proteins on Mb-scale *Caulobacter* chromosome compaction, we performed Bac-ATAC during exposure of *Caulobacter* cells to novobiocin, an inhibitor of both DNA gyrase and TopoIV (**SI Methods 4, SI Methods 13**). Bac-ATAC revealed that Mb-scale accessibility patterns of the *Caulobacter* chromosome changed significantly during novobiocin treatment, giving rise to 25 new or altered CAD boundaries (**Figure 2, Table 3**).

**Figure 2.**
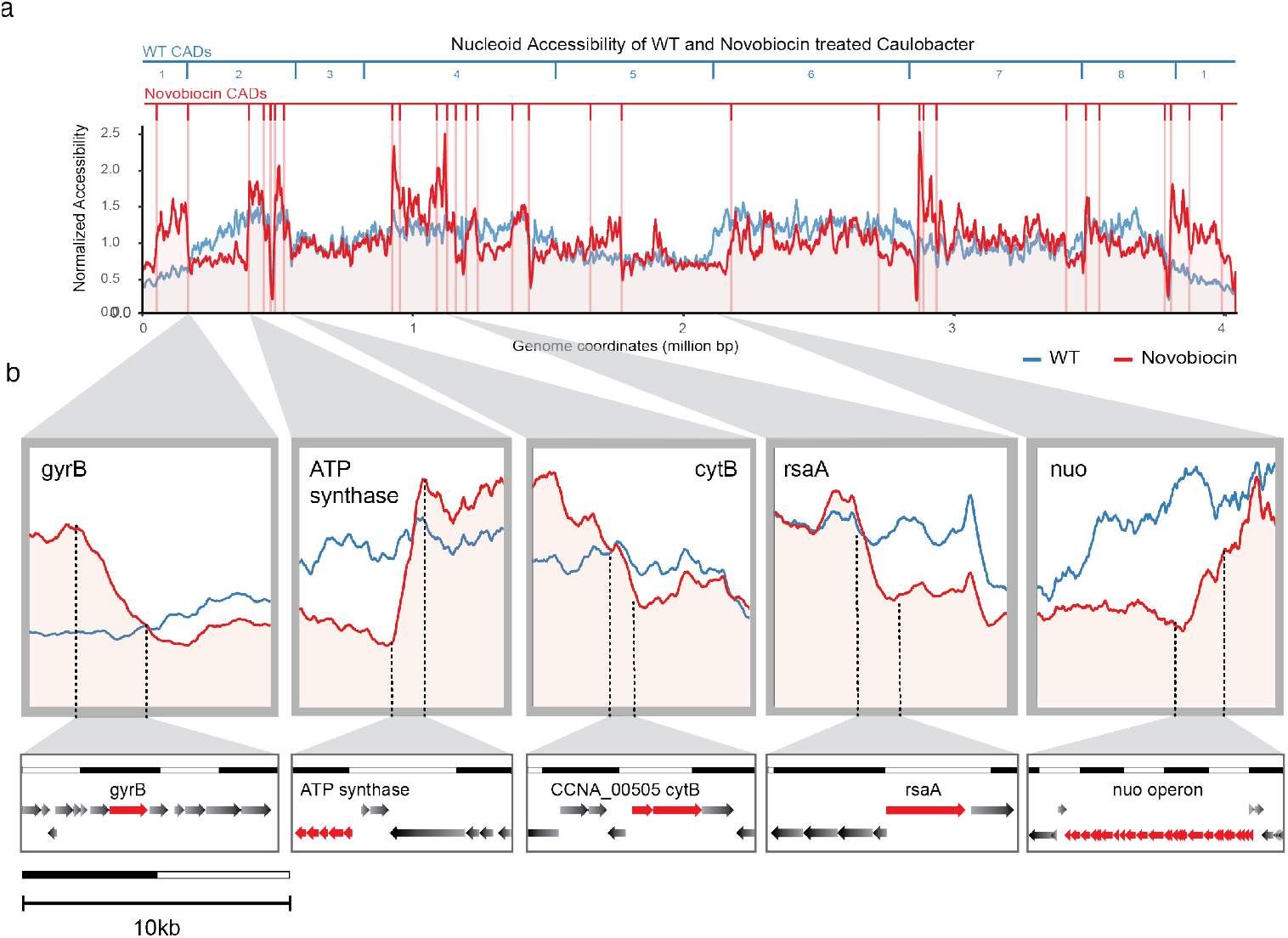
Bac-ATAC data for *Caulobacter* treated with novobiocin. **a**. Linear plot of Bac-ATAC data for novobiocin-treated cells (red) overlaid with WT data (blue). Red lines demarcate the Chromosomal Accessibility Domain boundaries found during novobiocin treatment (i.e., change-points in Bac-ATAC accessibility along the chromosome). WT CAD boundaries are enumerated along the top, in blue, for reference. **b**. Close-up images (red line), during gyrase inhibition, the regions of the chromosome overlapping these genes (or directly near these genes) form new accessibility change-points (i.e., at the depicted ATP synthase operon, *cytB* operon, *rsaA*, and *nuo* operon), or modify existing accessibility change-points (i.e. at gyrB). These accessibility change-points are identified as novobiocin-specific CAD boundaries and are further detailed in **Table 3**. Two of the longest Caulobacter operons encoding nuo (NADH-quinone oxidoreductase chain) genes as well as the operon containing genes CCNA_01984 to CCNA_01998, are oriented in the same direction on the chromosome and lie adjacent, near the terminus. Very long transcripts are somewhat associated with CID boundaries (15), and indeed this locus forms a CID boundary. Under normal growth conditions, this region is not a Chromosomal Accessibility Domain boundary. However, as seen in the final zoom-in panel, it emerges as a CAD boundary in novobiocin-treated cells. The *nuo* operon is the longest in *Caulobacter*, and extends beyond the region depicted in the rightmost panel.

Many of the new or altered CAD boundaries are located at highly expressed genes (**Table 3**; **SI Note 2**). For example, the *gyrB* encoding DNA gyrase, which is highly induced by novobiocin, forms an altered CAD boundary in this condition (**Figure 2b, Supplementary Data Set 1**). The novobiocin-induced CAD boundaries are also enriched for boundaries of the previously discovered Chromosomal Interaction Domain (CID) boundaries (4, 15); ten out of the 23 CID boundaries are recovered as new Bac-ATAC CAD boundaries in novobiocin treated cells (**Table 3, SI Note 3)**. Moreover, at the majority of all CID boundaries (19/23), novobiocin treatment created a pattern: the CID boundaries overlapping highly expressed or long genes in rich media became the locations of newly formed or altered CAD boundaries, while those associated with short or lowly expressed operons did not (**Table 4**). **Figure 2b** shows four CID boundaries that contain very long or highly expressed genes (the ATP synthase, *cytB, rsaA*, and *nuo* genes). In WT conditions, these four regions entirely lack CAD boundaries. During gyrase inhibition, however, these regions become accessibility change-points and can be identified as novobiocin-specific CAD boundaries. Other CID boundaries, listed in **Table 4**, follow this pattern as well.

**Table 4:**
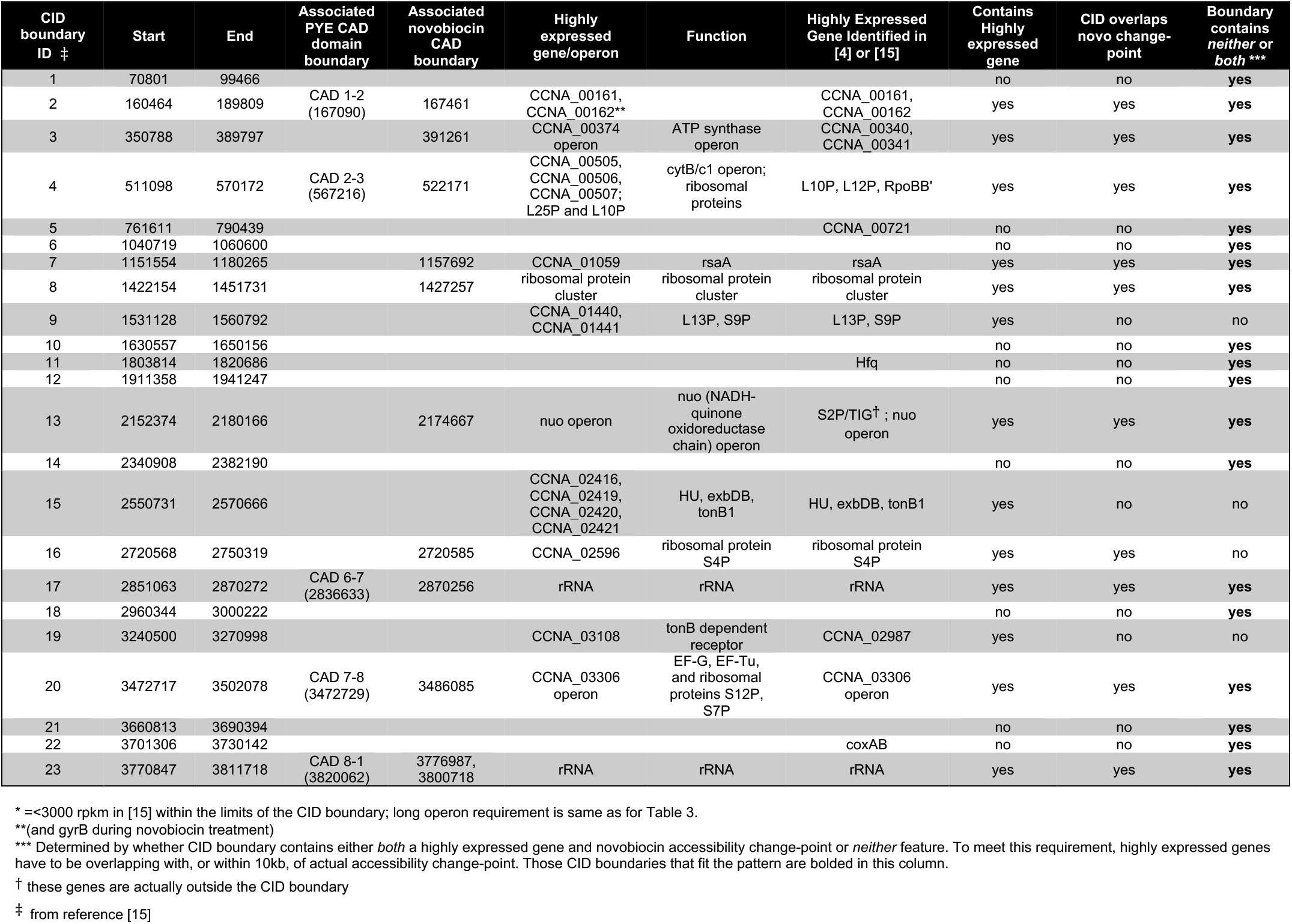
List of CIDs and their relationships to CAD boundaries and highly expressed genes in PYE, PYE + novobiocin

It is noteworthy that CID boundaries associated with highly expressed genes become CAD boundaries – but only under conditions where gyrase is inhibited. We propose that the aberrant CAD boundaries formed under conditions of gyrase inhibition are caused by un-checked transcriptional forces. Transcriptional elongation is known to increase positive superhelical density downstream of the transcription bubble, while increasing negative supercoiling upstream. When gyrase and TopoIV activity is inhibited during novobiocin treatment, increasing superhelical density overwinds the DNA downstream of the transcription bubble, whereas the upstream superhelical density is homeostatically kept close to levels reflecting WT conditions due to the action of other topoisomerases (16). We predicted that during novobiocin treatment, highly expressed and long operons associated with CID boundaries would cause extreme overwinding – and therefore increased compaction – of the DNA downstream of those transcribed regions and would subsequently lead to measurable decrease in accessibility of those regions. In agreement with this prediction, we observed that during novobiocin treatment, all accessibility change-points associated with highly expressed genes follow the same pattern: the direction of transcription matches the direction of decrease in accessibility (**Table 3**).

DNA gyrase is thought to maintain homeostatic levels of local DNA compaction throughout the chromosome, in large part by adding negative supercoils to the overwound DNA lying directly downstream transcription bubbles and replication forks. Here, we have provided evidence that gyrase activity also counterbalances the effects of transcriptional forces to maintain Mb-scale Chromosomal Accessibility Domain structure.

### Loss of GapR leads to loss of CAD boundaries

It has been shown previously that the essential nucleoid-associated protein (NAP) GapR binds globally to the *Caulobacter* chromosome *in vivo* (17). A structural study of the GapR-DNA interaction showed that GapR binds to the 3’ends of highly transcribed genes as a tetramer encircling the over-twisted DNA, thereby stimulating gyrase to relax positive supercoiling (18). Although GapR does not regulate gene expression, it is required for both the initiation and elongation of chromosome replication in *Caulobacter* (18, 19). Despite these findings, there is evidence that GapR performs additional functions in *Caulobacter*. Pleiotropic suppressors of a *gapR* deletion in many different genes not specifically related to DNA replication rescue growth and viability of *Caulobacter* (19). Furthermore, loss of GapR leads to errors in fidelity of timing of segregation initiation in *Caulobacter*, even in suppressors of *gapR* deletions that display WT initiation and elongation of replication by flow cytometry (19, 20) (**SI Note 4**). In *E. coli*, NAPs that play a role in the fidelity of segregation initiation also control chromosomal macrodomain structuring (21). In light of these observations, we asked if GapR contributes to CAD structure.

We obtained a suppressor of a *gapR* deletion that displays WT cell morphology (**SI Methods 5, SI Figure 5**). In the absence of GapR, Bac-ATAC revealed changes in the compaction pattern of the chromosome. Specifically, the distinct CAD boundaries observed in WT cells disappeared, leading to a smoothening of compaction levels over the chromosome (**Figure 3a; c**). To quantify this effect, we measured the average magnitude of the gradient of chromosome accessibility for the WT and mutant strains. This metric quantifies the integrated amount of change in chromosome accessibility occurring along each Bac-ATAC profile (**SI Methods 6)** (examples of scored accessibility patterns are shown in **Figure 3b**). The WT data score exceeded that of the *gapR* deletion data by a factor of three (**Figure 3c**), confirming that in the absence of GapR, the pattern of changes in nucleoid compaction along the chromosome smoothen out. While the changes in compaction along the chromosome are more gradual in the *gapR* deletion strain, the accessibility of the chromosome is not entirely uniform; we also observed lower compaction of the origin and terminus regions relative to the rest of the chromosome in the absence of GapR (**Figure 3a**), possibly as a result of cell shape (**SI Note 6**). Overall, we conclude that GapR activity structures CADs by controlling the formation of CAD boundaries.

**Figure 3:**
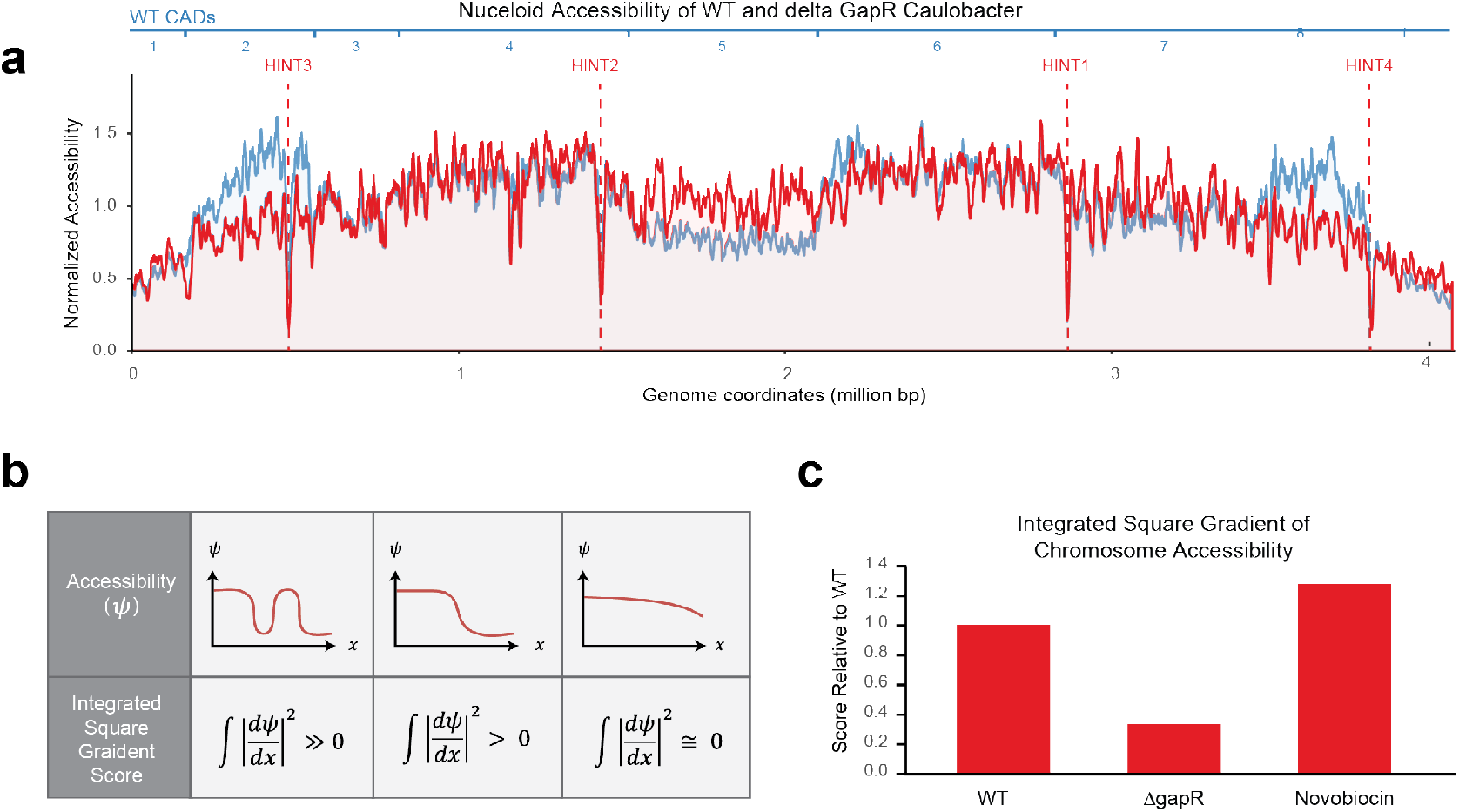
Bac-ATAC data for *Caulobacter* lacking GapR. **a**. Polar plot of Bac-ATAC data for *gapR* deletion suppressor mutants (red) overlaid with WT data (blue). WT CADs are enumerated on the outermost ring, in blue, for reference. **b**. Schematic for demonstrating how the Integrated Square Gradient describes the Bac-ATAC data. In the plots within this legend, *Ψ* represents accessibility in arbitrary units, and the X-axis represents base pair coordinate. The left-most and middle example curves represent example Bac-ATAC accessibilities that share identical standard deviations in accessibility. However, the leftmost example contains more change-points and would consequently have the greatest Integrated Square Gradient score (symbolized mathematically along the bottom row of the chart). In contrast, the rightmost curve represents example Bac-ATAC accessibility with the smallest Integrated Square Gradient; while like the middle example it crosses its average only once, it also has a smaller standard deviation. **c**. Plot of integrated square gradient metrics for the *Caulobacter* Bac-ATAC data in three conditions. This metric quantifies the extent to which the data tends to cross its own average (see **SI Methods 6**). The Y-axis represents the Integrated Square Gradient calculated relative to WT, which rests at value 1. The *gapR* deletion data shows a lower score and consequently has fewer change-points. The data from novobiocin-treated cells, on the other hand, has a higher score and contains more visible change-points.

To determine the effects of the absence of another NAP on CAD structure, we repeated Bac-ATAC in a strain of *Caulobacter* that lacked the Structural Maintenance of Chromosomes (SMC) protein. SMC has been shown to bind to and condense DNA globally in bacteria. In this strain, the CAD structure was unperturbed compared to WT cells (**SI Figure 6**).

### The GapR nucleoid-associated protein forms CAD boundaries, possibly by acting as a topological insulator

There are two main kinds of superhelical density that compact the bacterial nucleoid: non-diffusible superhelical density introduced by certain NAPs that bind, bridge, and bend DNA; and diffusible superhelical density introduced by transcription, replication, and topoisomerases. Topological insulators act as natural barriers to this diffusion, causing uneven accumulation of superhelical density in different parts of the chromosome (22, 23). It has been suggested that CIDs form topological domains of the Caulobacter chromosome, but evidence for this is limited (**SI Note 7**). We propose that CADs are true topologically insulated domains whose differential compaction levels result from their superhelical densities. We furthermore propose that the GapR NAP forms CAD boundaries by acting as a topological insulator, restricting the free diffusion of superhelical density between domains.

We have shown that CADs have characteristic average compaction levels. The most prominent line of evidence that CAD compaction is influenced by diffusive superhelical density is the observation that inhibition of DNA gyrase – a main effector of DNA superhelical density - causes drastic changes in Bac-ATAC signatures (**Figure 2**). Moreover, the largest changes in compaction in the absence of gyrase activity occur near highly expressed genes, where high levels of transcription introduce large amounts of diffusible superhelical density into the chromosome (**Table 3**). In contrast, deletion of the NAP SMC, which introduces non-diffusible superhelical density into the chromosome, did not lead to broad change in CAD structuring (**SI Figure 6**).

A topological insulator of CADs would i) form boundaries of differential compaction by ii) binding to CAD boundaries where it would iii) perform either DNA bridging or DNA wrapping to ultimately prevent free rotation of DNA about its central axis (2, 24). We propose that the main topological insulator of CADs is GapR. First, GapR forms boundaries of differential compaction, as its loss leads to smoothening out of superhelical density throughout the chromosome. Next, GapR binds to each CAD boundary in PYE (ie within ∼10kb) (**SI Table 8**) (17). Finally, there is evidence that GapR can prevent free rotation of DNA about its central axis. Using Atomic Force Microscopy, *in vitro* evidence has been obtained that GapR is capable of bridging DNA (25). Furthermore, Guo *et al*. recently demonstrated that GapR binds and constrains positive supercoils in DNA *in vitro*; if GapR can restrict loss of positive twist or writhe from DNA, it may also prevent diffusion of such structural motifs (24). We provide evidence that GapR performs this function even during novobiocin treatment (**SI Note 9)**.

Previous models have highlighted the role of GapR in synergistically activating DNA gyrase at the 3’-ends of genes *in vivo*, suggesting that gyrase function may be epistatic to GapR with respect to GapR’s effects on chromosome structure (18). In our model of the nucleoid, GapR and gyrase instead collaborate to structure CADs through distinct roles. On one hand, at many of the highly expressed genes that form CID boundaries, gyrase activity prevents the formation of CAD boundaries by evening out the local superhelical density gradients that would otherwise form due to transcriptional overwinding of DNA, possibly aided by its activator GapR. GapR binding, on the other hand, actually helps to form CAD boundaries by introducing topological boundaries that delineate Mb-scale regions of differential average compaction levels (see **Graphical Abstract**). Analysis of autocorrelation of Bac-ATAC data shows that these two proteins also have distinct effects on DNA structure on shorter (∼150 bp) scales, providing further evidence that gyrase is not completely epistatic to GapR with respect to its effects on chromosome structure (**SI Note 10)**.

Suppressors of *gapR* deletion display infidelity of chromosomal segregation initiation. We propose that the Mb-scale CAD structure imposed by GapR could be functionally important for fidelity of chromosome segregation initiation in *Caulobacter*, as is the case for the macrodomain structuring NAP MatP in *E. coli (21)*.

### A ribosome-associated HINT vanishes during phage infection

The lytic *Caulobacter* phage ϕCr30 harbors its own copy of *gapR* (17). We performed qPCR experiments on infected cells and observed that the phage *gapR* ortholog is expressed during infection of *Caulobacter* (**SI Figure 7a**). To determine the effect of infection on *Caulobacter* chromosome structure, we performed Bac-ATAC at twelve minutes after infection of *Caulobacter* swarmer cells with ϕCr30 (see **Materials and Methods 3, SI Methods 7**). Infection led to a drastic relative increase in the accessibility of the inaccessible (HINT) region formed by *Caulobacter*’s largest ribosomal protein operon (**Figure 4a, b, c**). This difference is shown quantitatively in **Figure 4d**. The Bac-ATAC profile of the rest of the chromosome remained similar to the no-infection control (**Figure 4d**).

**Figure 4.**
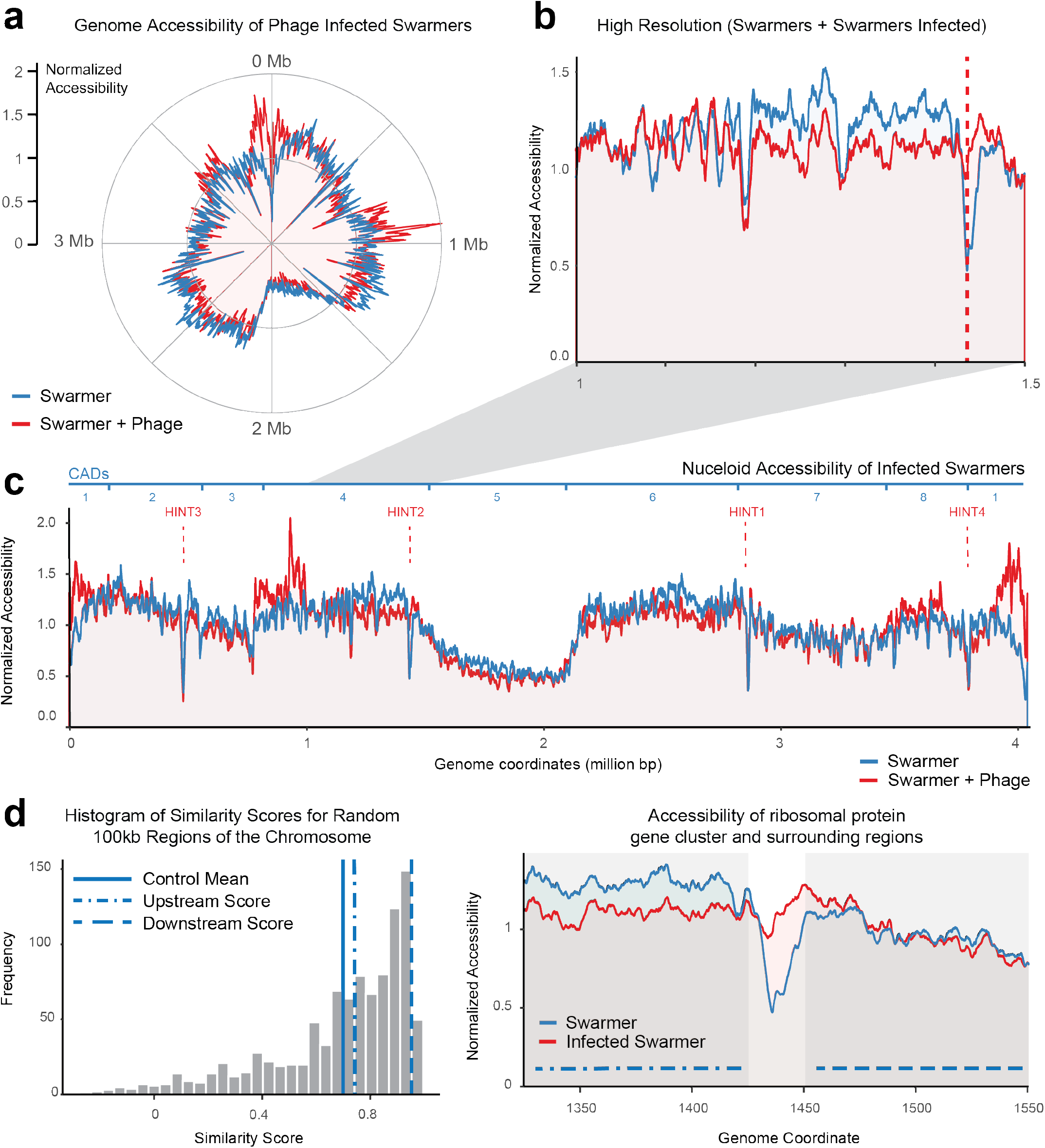
Bac-ATAC during infection of swarmer cells by the lytic phage ΦCr30. For polar and Cartesian plots of Bac-ATAC data, data was obtained for plotting as described in **SI Methods K. a**. Polar plot of Bac-ATAC data for *Caulobacter* swarmer cells. Swarmer cell data is blue; data for swarmers after incubation for twelve minutes with ΦCr30 phage is red. Polar coordinate: base pair coordinate, with origin at 12 o’clock. Bp coordinate increases in the clockwise direction. Radial coordinate: Chromosome accessibility determined from Bac-ATAC data was obtained for plotting as described in **SI Methods K. b**. Zoom-in of ribosomal protein cluster region during infection. This region is a highly inaccessible transcribed region in swarmers (represented by red dotted line) but becomes accessible during infection. Although there are subtle differences in accessibility between the infected and uninfected swarmer nucleoids, the most prevalent difference is at this locus. **c**. Cartesian plot of Bac-ATAC data for *Caulobacter* swarmer cells. Coloring is as in (**a**). HINTs are labeled with dotted red lines. WT CAD boundaries are enumerated along the top, in blue, for reference. **d. Left**: Histogram showing similarity scores for 1,000 random 100 kb chromosomal regions. The similarity score quantifies the extent to which the patterns in accessibility co-vary between the uninfected swarmer and infected swarmer. See **SI Methods 9** for more details. The mean for these control regions is represented by the solid blue line. **Right**: Another view of the ribosomal protein operon and its surrounding 100kb regions. The similarity score was also measured for the 100 kb regions to either side of the ribosomal protein operon (the ribosomal protein operon is shown as the lighter region in Cartesian plot of Bac-ATAC data). Accessibility scores for the upstream (dotdashed line) and downstream (dashed line) regions were both above average; these two scores are represented as the corresponding lines in the histogram. This implies that the accessibility of the nucleoid to either side of the ribosomal protein cluster region is similar between infected and uninfected swarmer cells.

We measured mRNA levels of the inaccessible ribosomal proteins spanning this locus during infection and discovered that the changes in expression displayed by these genes were mild and typical of other, distal ribosomal protein genes, none of which display this abrupt change in accessibility (**SI Figure 7b**). We conclude that changes in transcription at this locus are not likely the cause of the change in accessibility of this region. This implies that high transcriptional extent of this region is not the primary driver of its inaccessibility.

### Formation of highly inaccessible transcribed regions (HINTS) may reflect unique and functional physical properties of specific DNA regions

The *Caulobacter* chromosome is punctuated with <u>h</u>ighly <u>in</u>accessible transcribed regions (HINTs). A simple explanation is that extremely high transcription at these regions makes them naturally inaccessible to transposase. For example, transcriptional machinery may still contribute to blockage of transposase access to exceptionally highly transcribed HINT regions. However, many highly transcribed regions of the chromosome do not share the Bac-ATAC signature of HINTs (**Figure 1g, SI Figure 2**). Furthermore, we have shown that the control of the accessibility of the ribosomal protein HINT cannot be explained simply by its high transcriptional extent (**Figure 4, SI Figure 7b**).

Instead, the function of these regions may explain their inaccessibility. Most of the HINTs in *Caulobacter* are ribosome-associated in function (we discuss the others in **SI Note 11 and SI Note 12**). We also discovered ribosome-associated HINTs in *E. coli* (**SI Figure 3**). Importantly, these precise regions in *E. coli* have been shown to co-localize (26). Co-localization and interaction of these DNA segments within the nucleoid might also explain HINT formation; such organization could obscure the DNA from transposase. We propose that *Caulobacter* HINT regions may also co-localize. Evidence from multiple studies supports a recent proposal that co-localizing ribosomal DNA loci in *E. coli* form a bacterial nucleolus (27). This argument is based on the similarity between very highly transcribed DNA regions in bacteria and the eukaryotic nucleolus, which not only co-localizes, but which also phase separates (28). The precise mechanisms through which *Caulobacter* HINTs are formed remain to be discovered.

## Acknowledgements

Thanks to Dante Ricci for intellectual contributions; to Emma J. Chory for plentiful support and input, and for completing and designing many illustrations; and to Alicia Schep for contributing preliminary *E. coli* sequencing data and advice during early adaptation of the ATAC-seq method to bacterial cells. We thank for the Feig lab for generously sharing their chromosome structure models. Thanks to Stanford Center for Genomics and Personalized Medicine for completing whole genome sequencing, and to PAN for completing Bac-ATAC library sequencing. We acknowledge Jiarui Wang for performing both growth curves and imaging of strains.

## Materials and Methods

### Bac-ATAC protocol

#### Overall protocol

The Bac-ATAC protocol includes an adaptation of the ATAC-Seq protocol (8) to bacterial cells and an optimized reverse-crosslinking method (29). Bac-ATAC consists of five steps: fixation, permeabilization, transposition, reverse cross linking, and library preparation.

#### Fixation step

1-4 mL of culture at OD660 of 0.1-0.2 were collected and fixed in PBS pH 7.4 with 1% formaldehyde: (i) Make 1% formaldehyde in PBS pH7.4 on the day of the experiment. Keep formaldehyde solution on ice. (ii) Spin cells, then resuspend pellet in 1 mL ice cold PBS and incubate for five minutes at room temperature (RT)e; Hand-nutate tube for 30 seconds at the beginning and the end of the five minutes incubation. (iii) incubate cells for 15 minutes on ice. (iv) Spin crosslinked cells, discard supernatant. (v) Neutralize by repeating twice: resuspend and incubate for five minutes at RT with 1 mL 125 nM glycine. (vi) Collect cells, and *either* resuspend in 1 mL of PBS and keep on ice until freezing at −80 °C for no more than two weeks or continue to the permeabilization step. When thawing from freezer, thaw and spin cells before continuing with the permeabilization step.

#### Permeabilization step

(i) Wash cells gently in 400 uL 0.01 M Tris HCl pH 8 and then in 400 uL 0.033 M Tris HCl, 20% sucrose. Perform both spins at maximum speed for one minute. (ii) Resuspend cells in 400uL 0.033 M Tris HCl, 20% sucrose with 8 ul of 50 mM EDTA and 8 ul of 50 mg/ml lysozyme and incubated at 30 °C for five minutes, shaking at 400rpm. Prior to this step, keep lysozyme solutions on ice.

#### Transposition step

(i) Centrifuge 10% of the permeabilized cells for five minutes at 5000rpm. (ii) Wash in 800 uL of 0.033 M Tris HCl, 20% sucrose. (iii) Resuspend in transposition mix. Transposition mix contains 12.5 uL 2x TD buffer, 11.25 uL of 0.033 M Tris HCl, 20% sucrose and 1.25 uL Illumina Tn5 transposase (buffer, enzyme from Nextera XT kit). (iv) Incubate cells and transposase for 30 minutes at 37°C. If using a thermocycler, do not use heated lid. Hold completed reactions at 4°C.

#### Reverse crosslinking step

Reverse crosslinks by adding 200 uL of a mixture of 50 mM Tris–Cl, 1 mM EDTA, 1% SDS, 0.2 M NaCl, 5 ng/ml proteinase K to the transposition reaction and incubating at 65 °C for 14-17 hours, while shaking at 400rpm.

#### Library Preparation step

DNA was amplified following (8). Reactions were cleaned using the DNA Clean and Concentrator kit from Zymo. Libraries quality was measured using Bioanalyzer (HS run). Libraries were pooled and sequenced on a MiSeq using the 2×75 cycles.

### Analysis of Bac-ATAC sequencing data

#### Mapping transposon insertion points

Fastq files were trimmed using Flexbar FASTQ read trimmer, mapped to the NA1000 genome using Bowtie 2 using parameter –X2000. Option –m1 was not chosen, specifically so that rDNAs, which appear in identical copies in *Caulobacter*, could still be mapped (arbitrarily). Reads were then deduplicated using Picard Deduplicator. Unmapped reads were discarded, and the first mapped base pair of every remaining read was compiled to yield base-by-base insertion frequency counts for the entire chromosome. First mapped base coordinates were extracted from bam files using samtools command:

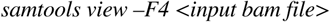

followed by implementation of a custom python code that extracts the first chromosomal base pair coordinate of each mapped read. This yields the precise location at which the transposase inserted donor DNA into the chromosome *in vivo*. Total insertion frequencies for each base pair coordinate of the chromosome were then compiled to generate raw Bac-ATAC data (**SI Figure 4, SI Table 4**).

#### Count normalization

The *in vivo* mixed population data sets were normalized to correct for copy number variations due to replication fork movements. To normalize, we generated corresponding *ex vivo* data sets by sequencing tagmented genomic DNA (gDNA). Both datasets were smoothened over 10kb windows. We then divided the smoothened *in vivo* data by smoothened *ex vivo* data.

For WT mixed population and *gapR* deletion strain MMDR503, data was compared to gDNA taken from the same strains grown in identical conditions to the respective experimental samples. For novobiocin-treated samples, the WT gDNA data was used for normalization. For swarmer and infected swarmer samples, normalization was not required for swarmer samples, as no replication occurs in these cells.

#### Collecting ΦCr30 infected swarmer cell samples for ATAC-seq

Swarmer cells are the only cell type that gets infected by ΦCr30 (10). An 8 mL sample of swarmer cells was obtained at the end of the synchrony protocol (below). Phage were added immediately as follows: 2.5 mL culture + 500 mL ΦCr30 phage lysate. The titer of this lysate was ∼1×10^9^ pfu/mL, so the Multiplicity of Infection (MOI) was ∼ 9. A mixture of 2.5 mL culture + 500 mL PYE served as an uninfected control swarmer sample. Samples were taken at 3 min and 12 min post-infection. Samples were not taken later than 12 min after infection to avoid contamination with newly differentiated stalked cells.

#### Synchronization of the Caulobacter cell cycle

Small-scale synchrony was performed as in (17) with the following changes: (i) We synchronized 60 mL of WT *Caulobacter* grown in PYE to OD660 0.44, (ii) We used Ludox in place of Percoll, (iii) We washed the cells in cold PYE instead of M2, and (iv) Final swarmer cells were resuspended in pre-warmed PYE.

## Supplementary Information

### Supplementary Notes

#### 1. ATAC-seq probes DNA compaction

We hypothesized that ATAC-seq probes chromosome compaction when analyzed on 100kb and higher length scales. This argument only requires the assumption that the transposase explores all regions of the nucleoid evenly, all else held equal (i.e., that transposase does not simply avoid entire Mb-regions of the chromosome more than others). This assumption is not unreasonable for two reasons. First, we have removed sequence-bias in our analysis by comparing to gDNA. Second, we are comparing Mb-scale regions that have no true differences in sequence or base composition at this level of resolution.

In **SI Methods 3**, we performed simulated ATAC and showed that compaction and accessibility of chromosomal DNA are inversely proportional on both the 15kb and 100kb length scales. Here we explore that there is furthermore a general relationship between compaction and accessibility from a geometric point of view. Mathematically, we let “⇔” imply a specific mathematical relationship. Below we will support the following postulate:

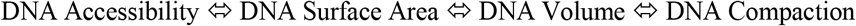

We would like to show that accessibility and compaction have an explicit geometric relationship *over lengths of DNA that have characteristic shapes*. Since Hi-C and other experiments have confirmed that the *Caulobacter* chromosome is toroidal and has a known and set orientation in space in the *Caulobacter* cell, we claim that regions of about 100kb or larger do in fact exhibit a predictable shape (1, 2). Because of this, the relationship Surface Area (SA) ⇔ Volume holds true for such segments of DNA, and is parameterized by the toroidal segment shape, where SA = 4 V/(R-r) and (R-r) is the difference in inner and outer radial lengths. Since compaction is essentially a form of DNA density, volume and compaction are inversely related by definition, with V = *b*/compaction (b is a proportionality constant). We note that from here on, we assume the DNA has toroid shape for sake of writing explicit equations. As long as the DNA has any predictable shape, the final result will be the same.

It remains to be shown that Accessibility ⇔ Surface Area. We assume that regions with larger *relative* surface area will then appear as being *relatively* more accessible to transposase. Mathematically, we only require that accessibility = *f*(r’) * (surface area) where r’ is radial distance from center of nucleoid. Since the transposase will access the cell from the outside, and reacts as it diffuses into the cell, w *f*(r’) will likely monotonically increase in r’ (although this is not a required assumption). We can now demonstrate that accessibility and compaction are also inversely related:

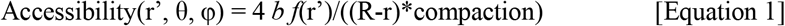

simplifies to

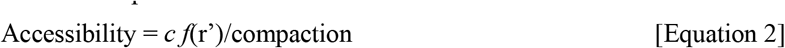

where *c* = 4 *b* /(R-r) (a constant). When comparing large (∼100kb) DNA regions to one another, we integrate over the coordinates (r’, θ, φ) the accessibility multiplied by DNA density for each set of coordinates. This would yield an approximation of relative expected insertion frequencies for each DNA region. If we assume that the distribution of DNA density in any two regions is nearly identical, then one part of the total integral, which appears as the following integral

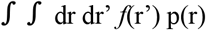

will be approximately equal for both regions. Alternatively, we could also assume that transposase reacts mostly with DNA at the outer surface (r’ =∼ R). Either way, after integrating we obtain the relation we have been looking for explicitly for comparing accessibilities of large regions:

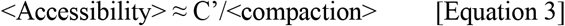

where “<>“ implies an average over the regions being compared. We have therefore shown using geometrical arguments that a region’s average accessibility and its average compaction should be inversely related and also inversely correlated. We can fill out our flow chart of mathematical relation as shown below:

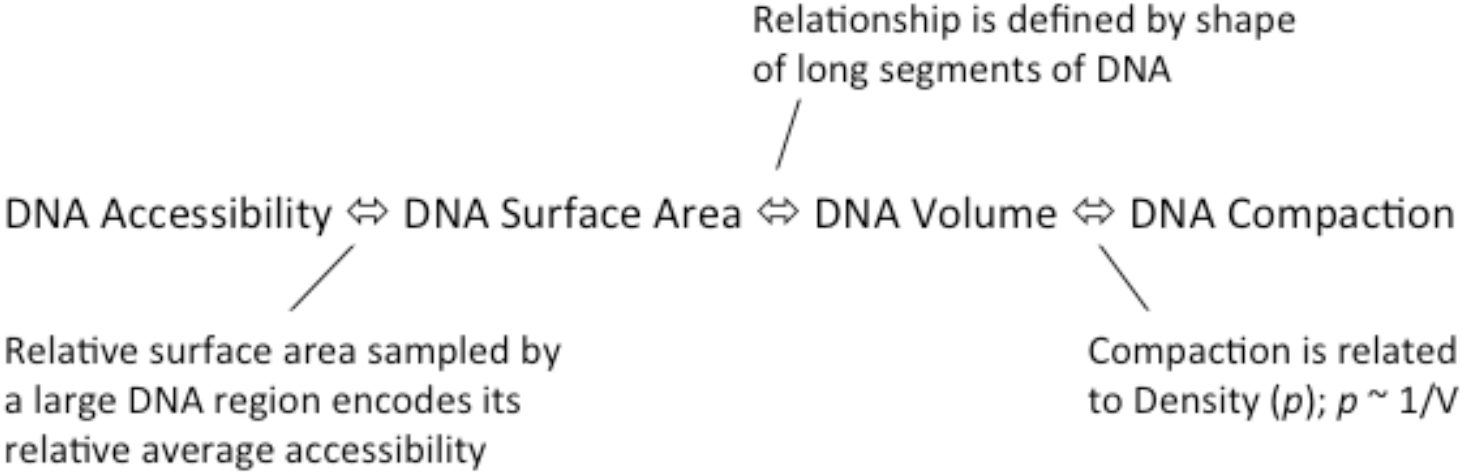

We also obtained experimental evidence in support of the relationship in **Equation 3**. Specifically, *Caulobacter* cells were exposed to sub-lethal doses of novobiocin in order to intentionally disturb chromosome compaction globally, and repeated Bac-ATAC. Novobiocin inhibits the activity of DNA gyrase, an enzyme that adds negative supercoils to DNA and thereby influences chromosome compaction in *Caulobacter*. As seen in **Main Text Figure 2**, inhibition of DNA gyrase globally disturbs the WT accessibility domains of the *Caulobacter* chromosome. Since DNA supercoiling is linked to DNA compaction and since changes in supercoiling caused by gyrase inhibition also led to changes in measured accessibility of different chromosomal regions, these results support the relationship expressed in **Equation 3**.

#### 2. RNA-seq during novobiocin treatment

We also performed RNA-seq during novobiocin treatment (**SI Table 6, Supplementary Data Set 1**). We discovered that five of the thirty total novobiocin change-points are located within genes and operons that are induced to high expression levels during novobiocin treatment. These genes included DNA gyrase subunits A and B and a drug efflux pump operon (**Table 3**, see **SI Note 3** for more details). The first zoom-in panel in **Main Text Figure 2b** illustrates how the chromosomal compaction pattern near WT CAD1-2 boundary, which is very near the *gyrB* gene, is inverted during novobiocin treatment. Whereas in WT conditions, accessibility increases sharply from left to right, during gyrase inhibition and *gyrB* activation, the pattern inverts and accessibility decreases sharply moving left to right (**Main Text Figure 2b**). In addition, more subtle change-points throughout the chromosome are associated with highly expressed genes during novobiocin treatment (**SI Figure 10**).

#### 3. Changes to chromosome accessibility during novobiocin-treatment to inhibit gyrase

Many of the change-points in novobiocin-treated cells are at highly expressed genes. The single bp-resolution of our method has revealed that the changes in accessibility are often centered over (or are very near to) specific, highly expressed transcripts (**Main Text Figure 2b**).

Indeed, 10/14 CIDs associated with highly expressed genes become CADs upon the inhibition of gyrase by novobiocin. An additional two (CIDs #9 and #15) form less significant CADs and so do not count towards this total (**Table 3**). In some cases, highly expressed genes may be less likely to form significant CAD boundaries because they are convergently transcribed with other highly expressed genes (i.e., at CID #15). This convergent transcription may obscure the ability of these regions to form change-points if, in order for change-points to form, superhelical density must build up asymmetrically near genes and operons that are highly transcribed in one direction in the absence of gyrase activity. Finally, there is a fundamental limit to our ability to accurately identify CAD boundaries using the Bac-ATAC method, since many factors likely affect chromosome accessibility and since true accessibility change-points may occur to varying degrees on different length scales.

**SI Table 6** lists the most highly expressed genes during novobiocin treatment and **SI Figure 10** shows these gene locations overlaid onto the novobiocin ATAC data. For complete RNAseq during novobiocin treatment data, see **Supplementary Data Set 1**.

#### 4. General notes on previously-identified GapR suppressor mutants

Many suppressors of *gapR* deletion have been reported and mapped to various regions (3, 4). These regions include ribosomal RNAs, regions upstream of tRNAs, and the RNA polymerase gene *rpoB (3)*. In this work we report another *gapR* suppressor mutation in the gene downstream from *gapR*, which we have named *gapS* (**SI Methods 5**). It is unlikely that these diverse suppressor mutations all function to rescue WT rates of replication elongation directly, implying that GapR may serve multiple functions in the *Caulobacter* cell cycle.

It has also been shown in reference (3) that the mutations in the origin of replication (strain 3880) and in the 23S rRNA (strain 3920) significantly ameliorate or rescue, respectively, replication initiation and elongation phenotypes of *gapR* deletion strains. Importantly, however, the suppressor mutation in strain 3920 rescues WT replication initiation and elongation but does not rescue other cell cycle errors. For example, strain 3920 displays segregation errors. This observation led us to ask whether GapR could contribute to fidelity of chromosome segregation, and whether it did so by controlling CAD structure.

#### 5. Accessibility of *rsaA* gene locus in a *gapR* deletion mutant

The absence of GapR also led to a notable decrease in the accessibility of the highly expressed *rsaA* gene. *rsaA* is the most highly expressed mRNA in *Caulobacter*, and codes for protein that composes the extracellular crystalline S-layer (5, 21).

#### 6. Notes on possible effects of cell shape on ATAC data in the *gapR* deletion mutant

Cell shape may cause the arms of the chromosome to appear more accessible; they are afforded more surface area due to the elongated shape of *Caulobacter* cells coupled with the ori-ter arrangement of the chromosome.

#### 7. CIDs as topological domains

It has been proposed that Chromosomal Interacting Domains (CIDs) represent topological domains in the *Caulobacter* nucleoid (5). The authors claim that CID boundaries tend to form near highly expressed genes, and propose that high transcription levels at these regions, through inhibition of supercoil diffusion or other mechanisms, lead to formation of topological boundaries. However, the evidence that CIDs behave as true topological domains is limited. First, upon close inspection, the connection between transcription of highly expressed genes and the formation of either CID or CAD boundaries is actually quite tenuous. Neither CID boundaries nor the Bac-ATAC-derived CAD domain boundaries are consistently near highly expressed genes or long operons (**Table 2, SI Table 2**). In addition, the physical properties like the superhelical density of individual CIDs has not previously been measured. Although CIDs are certainly interacting domains, their true topological properties (like their superhelical densities or compaction levels) cannot be determined with proximity ligation assays alone.

The lack of relationship between true topological domain boundaries and CID boundaries is actually consistent with our results, which demonstrate that CID boundaries – while demarcating DNA-DNA contact barriers – do not demarcate changes in other physical properties of DNA. For example, CID boundaries do not generally align with accessibility change-points (CAD boundaries) (**SI Figure 1**). As we argue in the **Main Text**, this is in part because the transcriptional forces that might otherwise form accessibility change-points (i.e., CAD boundaries) at CID boundaries are actually suppressed by DNA gyrase activity. A key example is *rsaA*. Under normal growth conditions, this region forms a CID boundary but does not form a CAD boundary. Strikingly, when we inhibited gyrase activity and repeated ATAC-seq, the majority (10/14) of CID boundaries that are found near highly expressed genes – including *rsaA* – are recovered as CAD boundaries (**Table 2, Main Text Figure 2b**).

#### 9. Evidence that GapR may function as a topological insulator even in novobiocin treated cells

It has been shown by chromatin immunoprecipiation (ChIP) that GapR binds within 10kb of all CAD boundaries found in novobiocin-treated cells except three (**SI Table 9**) (3). One of the CAD boundaries apparently not near GapR in this table is actually associated with GapR, but this association is obscured by the presence of a nearby HINT region (the ribosomal protein operon). This long region mostly binds GapR at its 3’ edge, further downstream from the computationally-determined CAD boundary.

For the remaining two novobiocin-condition CAD boundaries, it may be that GapR binding distal (∼30kb) from the boundary can still play a role in allowing for disproportionate accumulation of superhelical density downstream one side of the highly expressed operons associated with those boundaries. For example, one of these CAD boundaries, novo CAD #19, overlaps the *nuo* operon, which lies adjacent to another long operon (CCNA_01984 to CCNA_01998). GapR binds to the 3’ end of this second operon (according to data from reference (3)). In this case, GapR may appear to bind far away from the change-point center, but GapR may still help define this CAD boundary by acting as a topological insulator at the 3’ end of these very long operons (**Main Text Figure 2b**, fifth zoom-in panel).

#### 10. Autocorrelation analysis of raw Bac-ATAC data

We measured the autocorrelation of the raw insertion frequency data for WT cells, *gapR* deletion strain MMDR503, WT cells + novobiocin, and naked genomic DNA from WT cells (mixed population), as well as from swarmer cells from WT *Caulobacter*. The autocorrelation scores were normalized and plotted together versus lag (the lag is a measure of distance in base pairs between points that were compared for the calculation performed across the chromosome, with each lag value being assigned a different x-coordinate in the graph) (see **SI Figure 9**). Autocorrelation for intact WT mixed populations and swarmer cells, as well as *gapR* deletion strains, look identical, with continued periodicity. In contrast, novobiocin-treated cells have accessibility autocorrelation profiles similar to that of purified genomic DNA, which features no periodicity in high lag regime (more than about 30bp). The autocorrelation is cut off in “y” at 0 bp where it is arbitrarily large.

The autocorrelation at low lag may represent periodicity in the structure of DNA, where the helical turn is between 10-11 bp long. This is the distance at which we see the largest peaks in autocorrelation. A similar pattern of periodic enzymatic activity on DNA has been reported previously for DNaseI digestion (6). There is also periodicity in the frequencies of insertions at longer length scales. This implies that intact DNA has periodical topological patterns in intact cells that extend for longer distances (i.e., for at least 150 bp). The distance we measure may be limited, among other things, by the distribution of fragment sizes that we sequenced. We note that even though these patterns appear in the absence of GapR, they disappear in both extracted DNA and in cells in which gyrase is inhibited.

GapR and gyrase have distinct effects on other structural properties of the chromosome as well. Analysis of the autocorrelation functions of the raw Bac-ATAC data from WT mixed population and swarmer cells shows that the structure of the nucleoid is highly periodic, with periods on the scale of ∼10 to ∼150bp. We note that this autocorrelation pattern is missing for naked DNA (**SI Figure 9**). While the nucleoids of a *gapR* deletion strain demonstrated the same pattern as in WT cells, inhibition of gyrase creates autocorrelation pattern highly similar to that of naked DNA (**SI Figure 9**). This implies that DNA gyrase is at least partially responsible for the WT periodicity in DNA accessibility on short length scales. It furthermore means that even in the absence of GapR, DNA gyrase activity still contributes to maintenance of WT DNA structure on a local level.

#### 11. Notes on HINT detection by ATAC-seq

HINT regions do not arise as mapping errors, since non-zero transcript levels are carefully assigned even when non-unique sequences are mapped (albeit randomly) to the two rRNA loci rather than being discarded. These regions furthermore always obtain some insertions, implying that their access is not immeasurably low. HINT regions *can* be sequenced, since the ribosomal protein HINT appear as accessible during phage infection experiments where it accrues insertions (see **Main Text Figure 4b**). Additional evidence that these regions are not consequences of mapping errors lies in that fact that the ribosomal protein and prophage element-associated HINTs are not duplicated regions (the two rRNA loci are identical and could, without care, lead to mapping errors). Nevertheless, the ribosomal protein and prophage element-associated HINTs appear specifically inaccessible.

#### 12. Notes on properties of the *parS* locus

The *parS* centromere region is specifically inaccessible in swarmer cells (**Main Text Figure 1**). This region is the nucleation site for ParB protein, which ultimately helps guide one daughter chromosome to the opposite pole during chromosome segregation during the cell cycle. It was recently shown that ParB spreads over the precise ∼10kb stretch of DNA surrounding *parS* we identify as inaccessible by Bac-ATAC (20). GapR seems to delineate this highly inaccessible region, as it binds to either side of the centromere (7). We predict that in the swarmer cell, this region is specifically made inaccessible by its interactions with proteins involved in regulation of chromosome segregation including ParB.

### SI Methods

#### 1. Optimizing the Bac-ATAC Protocol

We monitored *Caulobacter* cells microscopically for lysis during the Bac-ATAC procedure, specifically at the stage where cells are permeabilized and incubated with transposase. We wished to minimize cell lysis because we were interested in monitoring physical states of the intact nucleoid with this method. Addition of sucrose during permeabilization significantly reduced lysis when cells were incubated with mock transposition mix (Nextera XT TB buffer at 1x concentration, but without transposase). Crosslinking cells, as was done during ATAC-seq (7), in combination with addition of sucrose, helped further mitigate cell lysis.

#### 2. Defining CADs and CAD boundaries

Smoothened, normalized mixed population data (**Main Text Figure 1**) was subjected to change-point analysis to detect the locations of abrupt changes in chromosome accessibility. Smoothening was performed using sliding window smoothening with window size = 100,000 bp. Then, change-points were found using MATLAB’s built in findchangepts function, with N = 9. One change-point that was originally found within 150kb of other change-points was eliminated for simplicity. **SI Figure 11** shows an alternative view of Chromosomal Accessibility Domain architecture of the *Caulobacter* chromosome where all nine original change-points are identified.

#### 3. Accessibility versus Compaction analysis

We analyzed an ensemble of high-resolution three-dimensional models of the single chromosome of swarmer cells (1) to study the relationship between *Caulobacter* chromosomal DNA accessibility and compaction. The swarmer chromosomal DNA was modeled as a closed worm-like chain of segments with an equilibrium length of 5 nm each representing 15 base pairs and a radius of 1 nm. An ensemble of 1026 structural models was clustered into 27 groups based on pairwise mutual similarity; Each cluster populated between 0.5 to 8 percent of the ensemble structures (1). The ensemble was shown to be consistent with the swarmer cell dimensions (8), Hi-C data (9), and intramolecular distances between 51 fluorescent-labeled DNA segments (10). We adapted FreeSASA (11) to calculate the accessibility of each structural model in the ensemble to a transposon molecule binding. FreeSASA provides an efficient implementation of the Lee and Richards algorithm for calculating solvent accessible surface area (SASA) (12). The radius of each chromosome structure bead was set to 3 nm and the radius of the rolling sphere was set to 2.5 nm based on the radius of gyration of the *E. coli* transposon structure (13).

To calculate the compaction of each structural model in the ensemble we calculate the number of neighboring beads in a 25 nm sphere centered at the query bead. For efficient calculation we used the scipy k-dimensional tree implementation (scipy.spatial.cKDTree). To determine the local relationship between accessibility and compaction we first averaged the accessibility and compaction scores in 15000bp windows and calculated the Pearson correlation between these two metrics. For all 27 clusters the Pearson correlation indicated anticorrelation with values lower than −0.95, suggesting that high accessibility correlated with low compaction and vice versa (**SI Figure 8**).

#### 4. Identification of CAD boundaries in novobiocin treated cells (data shown in Main Text Figure 2)

The protocol outlined in **SI Methods 2** was used to identify change-points in Bac-ATAC data from novobiocin-treated cells (**Main Text Figure 2**) with parameter N set to 30 and without an additional round of smoothening. See **SI Methods 13** for details of novobiocin treatment.

#### 5. *gapR* deletion suppressors used in this study

We had demonstrated that a deletion of gapR impairs growth (14). To study a viable deletion strain, we obtained a suppressor of a *gapR* deletion (strain MMDR 503). Initial deletion of gapR was performed using homologous recombination as described in reference (14). Rare *gapR* deletion candidate strains were isolated and confirmed to be true deletion strains. Phage ΦCr30 lysates of those strains were generated. One confirmed *gapR* deletion was transduced into WT *Caulobacter crescentus* by the standard protocol for generalized transduction of *Caulobacter* with ΦCr30 (as performed in (14)). After several days of incubating plates at room temperature, colonies formed. Colonies were purified, grown in liquid media, and frozen.

The specific deletion strain used here, MMDR 503, does not have significant morphological abnormalities compared to wild type *Caulobacter*, but grows slowly compared to WT cells (**SI Figure 5**).

Using whole genome sequencing of MMDR 503, we mapped the suppressor mutation to the gene downstream of *gapR*, which we name *gapS*. Sequencing also confirmed the in phase *gapR* deletion. The suppressor mutant (*gapS*\*) contains a single amino acid substitution (L76P) that may interrupt an alpha helix in this protein. This protein has some homology to toxin proteins but is not predicted to bind DNA. Although GapS is not essential (4), this is not surprising since it is barely translated in WT *Caulobacter*, with one of the lowest translational efficiencies in the whole genome (15). We furthermore have preliminary evidence that translation of *gapS* from the vanillate-inducible promoter/RBS in *Caulobacter* is toxic (data not shown). We predict that GapR/GapS may behave like a toxin-antitoxin system in *Caulobacter*.

#### 6. Quantification of the integrated amount of change in accessibility along the chromosome measured by Bac-ATAC

We calculated the average magnitude (square) of the gradient (first derivative with respect to bp coordinate) of the Bac-ATAC data. To do this, we binned the smoothened and normalized accessibility data by 100,000 bp, and calculated a fractional change in accessibility between adjacent bins. The sum of the square of these differences was added for each strain and compared. The final value quantifies the extent to which the accessibility varied over the whole chromosome between bins of many kilobases. The standard deviation metric does not capture this information, since it only measure the spread in the data without information about the frequency with which the data crosses its average.

The results are shown in **Main Text Figure 3c**. This metric, when calculated for the WT data, is three times that obtained for the *gapR* deletion suppressor strain. The data obtained for the novobiocin-treated cells had a score higher than that obtained for WT cells. This is consistent with the observation that this data displays more frequent changes in accessibility along the chromosome and is characterized by a greater number of CAD boundaries.

#### 7. Analysis of phage infected swarmer cells

##### Synchronization of the Caulobacter cell cycle

Small-scale synchrony was performed as in (14) with the following changes: 60 mL of cultures of WT *Caulobacter* grown in PYE to OD_660_ 0.44 was synchronized. Ludox was used in place of Percoll. All washes of cells with M2 were performed as washes in cold PYE. Final swarmer cells were resuspended in pre-warmed PYE.

##### ΦCr30 infection for ATAC-seq

WT *Caulobacter crescentus* NA1000 was grown in rich media (PYE) to midlog and swarmer cells were isolated by differential centrifugation (14). Swarmer cells are known to be the only cell type that gets infected by ΦCr30 (16). An 8 mL sample of swarmer cells in PYE rich media was obtained at the end of the synchrony protocol (the OD_660_ of this culture was 0.22 once the synchrony was complete). Phage were added immediately as follows: 2.5 mL culture + 500 mL ΦCr30 phage lysate. The titer of this lysate was ∼1×10^9^ pfu/mL, so the Multiplicity of Infection (MOI) was ∼ 9. A mixture of 2.5 mL culture + 500 mL PYE served as an uninfected control swarmer sample.

##### Obtaining ATAC-seq samples from infected cultures

Cultures were grown on a rotating wheel at 29**°**C, and samples were taken at 3 min and 12 min post-infection (matching the timing of phage infection qPCR experiments described below). Samples were not taken later than 12 min after infection to avoid contamination with newly differentiated stalked cells. ATAC-seq was performed on samples as described in **Methods**.

##### qPCR of phage infected cells

Cultures of *Caulobacter* NA1000 were inoculated from single colony of WT *Caulobacter* grown in PYE to OD_660_ = 0.6. Cultures were then diluted by mixing 2 mL of culture with 3 mL fresh PYE, and incubated for two hours. Midlog cultures were then exposed to phage or to sterile filtered DI water. For infection, cultures were mixed 5:1 with transducing lysate of ΦCr30 (titer ∼1×10^9^ pfu/mL as was used for the ATAC-seq experiments). MOI was again ∼9. A mixture of 500 mL culture + 100 mL sterile deionized H_2_O served as an uninfected control. The samples were swirled briefly and incubated at room temperature (without shaking or spinning). 1 mL samples were collected at each time point at the following times: just before introduction of phage, then at 3 min, 12 min, and 30 minutes after infection. Before freezing, each sample was vortexed to remove adsorbed phage, pulse-spun for 15 seconds, supernatant was removed, and pellets were flash frozen in liquid nitrogen.

##### qPCR Targets

This table lists the primers used for the qPCR experiments.

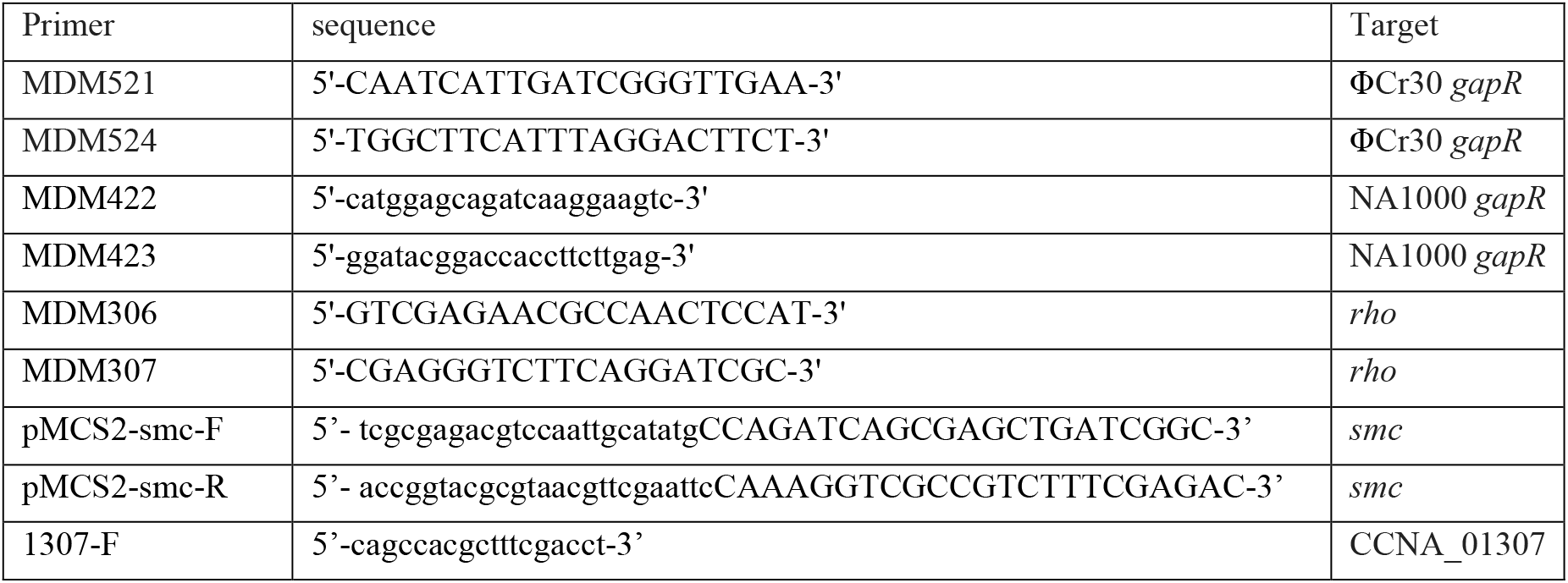
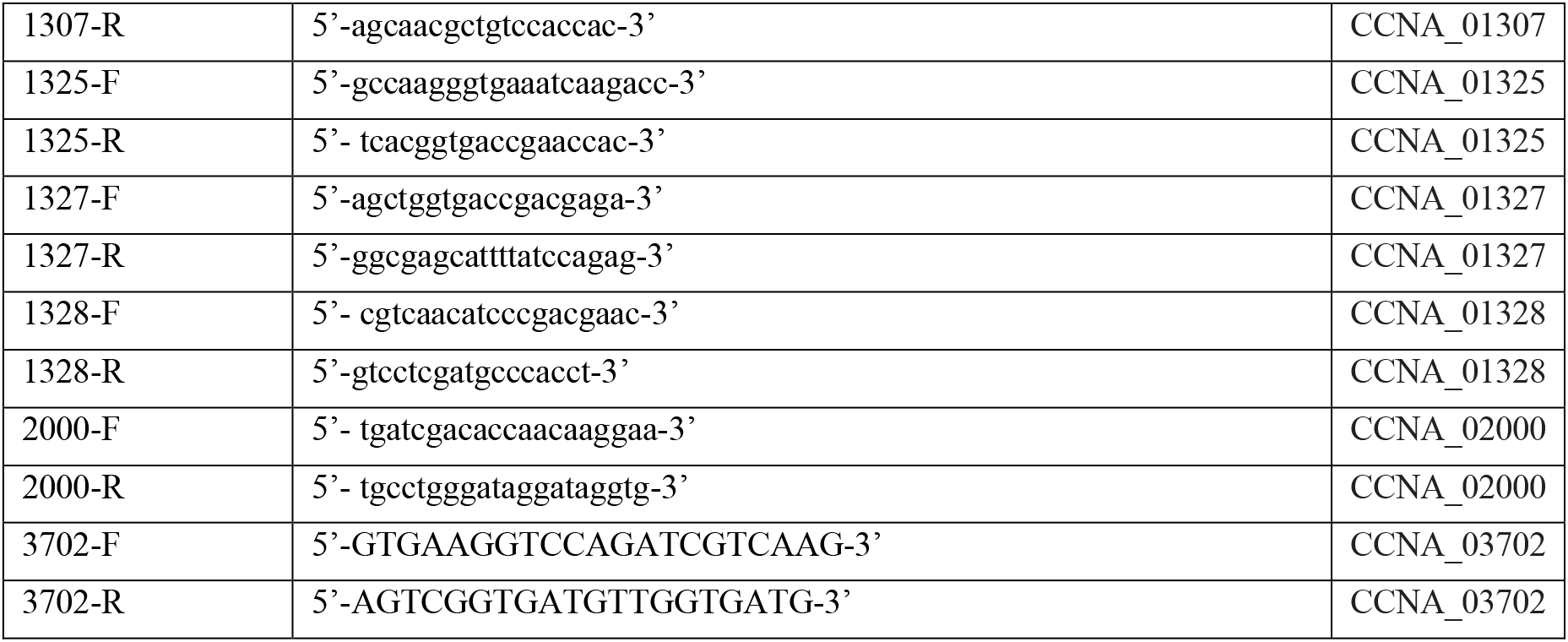

For phage infection-qPCR, primers were carefully tested for target specificity by amplifying on a mixture of genomic DNA from both NA1000 and ΦCr30 by melt-curve analysis.

The qPCR primers that amplified phage *gapR* and *Caulobacter gapR* cDNA were tested for target specificity by by melt-curve analysis, using a mixture of genomic DNA obtained from both WT NA1000 and the phage ΦCr30 as template. Only those primer pairs that produced a single, unique product in the presence of both phage *gapR* and *Caulobacter gapR* DNA templates by melt curve analysis were used in the qPCR experiments. RT-qPCR was performed as in (14). Also as in (14), primers specific to *rho* were used for amplification of the internal standard.

##### Choosing qPCR targets for phage infected swarmer cells

The genes tested were those that displayed specific changes in accessibility during infection of swarmer cells by ΦCr30 (**Main Text Figure 4**). The genes showing the greatest change (with the exception of one *rpoA* gene) are all those encoding ribosomal proteins within two very proximal genomic windows. These genes are separated by two genes of unrelated function, *secY* and *adk* (CCNA_01326 and CCNA_01327).

The first window in which we measured expression was [CCNA_01307 to CCNA_01323] including those boundary genes. The second window in which we measured expression stretches from [CCNA_01328 to CCNA_01332] again including boundary genes (see data in **SI Figure 7**).

#### 8. genomic DNA preparation

##### Genomic DNA Preparation

Genomic DNA was isolated from *C. crescentus* strain NA1000 using the Sigma GenElute Kit following the included protocol. The following adjustments were made: tagmentation steps for library prep followed genomic DNA extraction used either Ultrapure water or 10 mM Tris pH 7.5. No EDTA was used.

##### Phage genomic DNA purification

We followed the “PCI/SDS DNA extraction” protocol from PhagesDB (17).

#### 9. Calculating similarity in accessibility between experimental and control regions of swarmer cell nucleoids during infection

The similarity scores shown in **Main Text Figure 4d** were calculated by taking the dot product of the Bac-ATAC data from both the infected and uninfected swarmer cells. Smoothened accessibility data was standardized and then normalized, and a dot product was calculated. A score of 0 implies no correlation of accessibility patterns whereas 1 implies correlation. The control data was obtained by sampling 10,000 random windows, each 100kb in size, and calculating the same dot product. The data in the histogram in **Main Text Figure 4d** shows the distribution of similarity scores of these random regions.

#### 10. Whole genome sequencing

Whole genome sequencing was performed on genomic DNA extracted from strain MMD503. DNA was drop-dialyzed and resuspended in Ultrapure water to remove EDTA. Tagmentation was performed by the Stanford PAN facility using Nextera XT tagmentation kit followed by whole genome amplification. Libraries were QC’d by Bioanalyzer and Qubit, and multiplexed. Paired-end sequencing of multiplexed libraries was performed by Stanford Center for Genomics and Personalized Medicine facilities on an Illumina Hi-Seq.

#### 11. General notes on making Main Text Figures, SI Figures 1,**3**

Unless stated otherwise, plotted data was gathered from a mixed population of *Caulobacter* grown in PYE to midlog phase. Unless stated otherwise, data was normalized by dividing the given experimental data by a reference data set after both data sets were smoothened using sliding window averaging, with a window size of 10kb. Before the resulting smoothened, normalized data was plotted it was again normalized so that its average value is 1 for comparison to data obtained from other samples.

#### 12. Microscopy

Agarose pads were made from 1.5% agarose gel in M2G. Images were taken with Hamamatsu EMCCD and on a Leica DMi8 microscope. For images in **SI Figure 5**, exposure time was set to 150ms.

#### 13. RNA extraction for RNA-seq during novobiocin treatment

*Caulobacter* cultures of log-phase cells (OD_600_ = 0.15) were incubated with 50 ug/ml novobiocin, or water as a control, for 30 min at room temperature. The extracted RNA from each sample was stabilized by adding two volumes of RNAprotect Bacteria Reagent (Qiagen) follow by a standard total RNA extraction using RNeasy Mini Kit (Qiagen).

#### 14. RNA sequencing

The library preparation and RNA-seq were conducted by Novogene.

#### 15. RNA extraction for qPCR experiments, and qPCR protocol

These experiments were performed as in reference (14).

## Supplementary Figures

**SI Figure 1.**
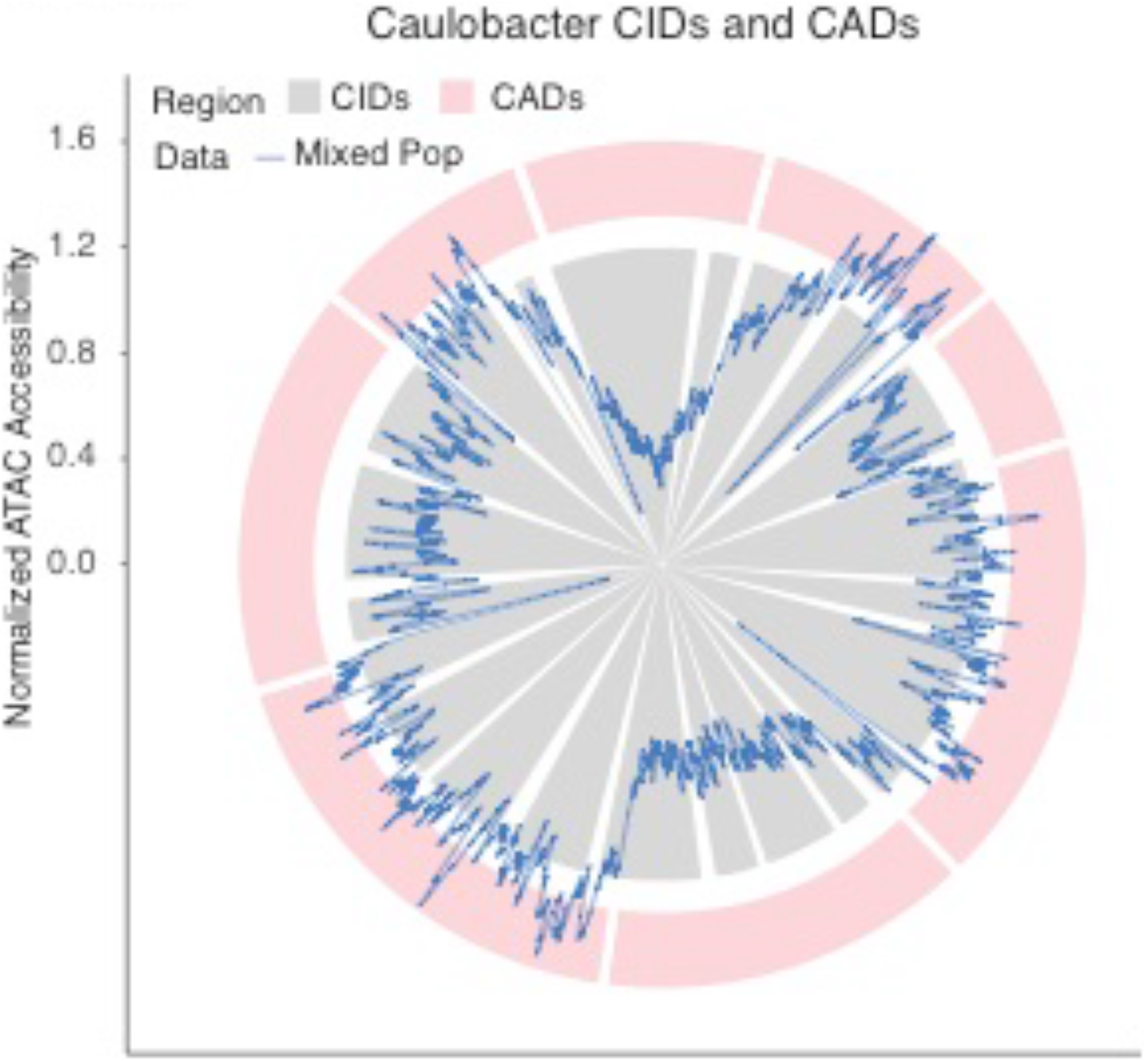
CIDs and CADs overlaid on the chromosome. CIDs (grey) overlaid with CADs (pink). Domain boundaries are in both cases highlighted by white gaps (CID boundaries are from reference (5); white gaps between CAD boundaries are exaggerated for visibility). Coordinates are the same as in **Main Text Figure 4** with bp coordinate along the angular coordinate and chromosome accessibility along the radial coordinate. Bac-ATAC data from WT cells (blue line) is replicated from **Main Text Figure 1**. We note that while CADs generally contain groups of CIDs, that CAD boundaries do not precisely overlap CID boundaries. CID boundaries and CAD boundaries overlap in 3/8 cases. CADs are generally composed of groups of CIDs, since all CADs are made up of at least two CIDs.

**SI Figure 2.**
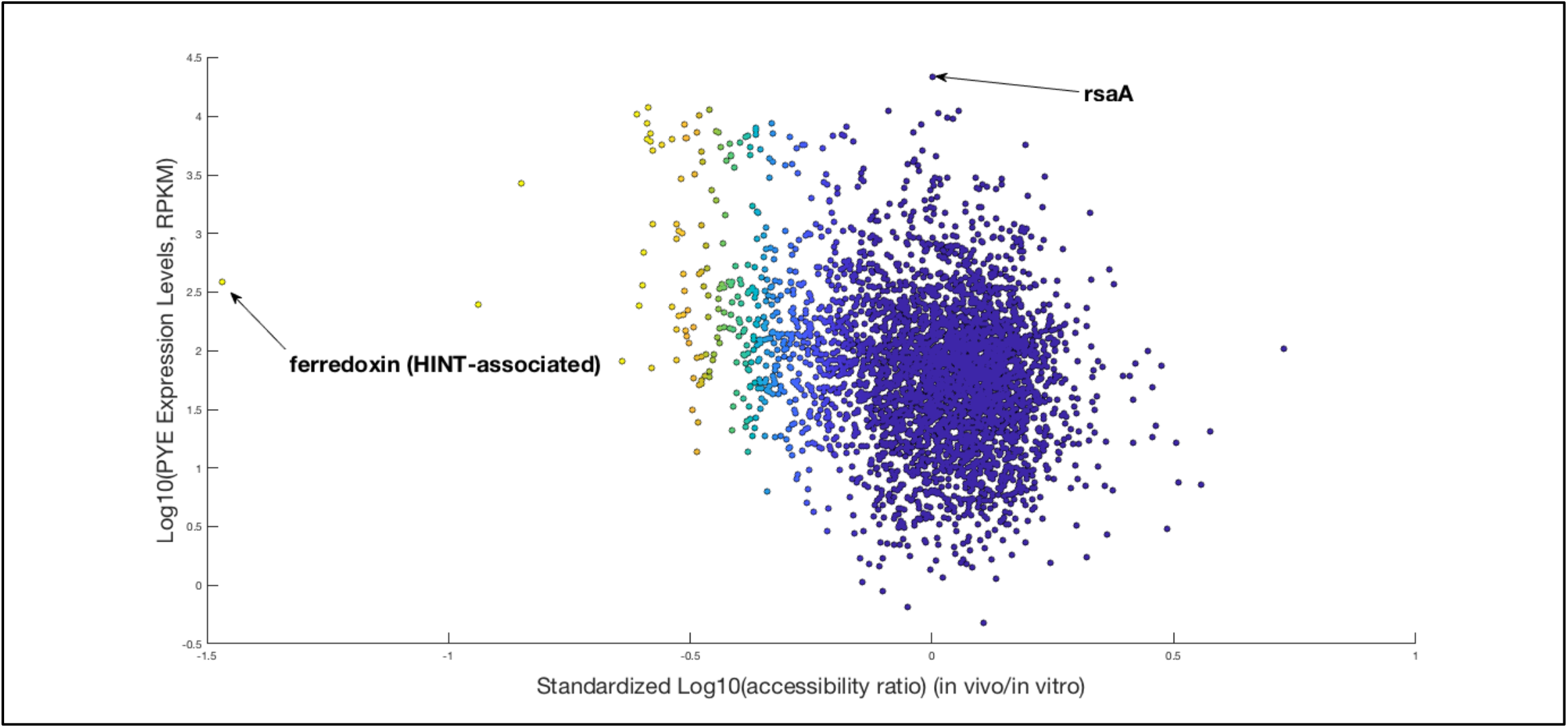
Scatterplot of log-log expression versus accessibility for all transcripts. Log-Log plot of Expression vs accessibility for all genes in *Caulobacter*. Color encodes a logarithmic scaling of the x-axis to highlight the very most highly inaccessible genes (yellow = much less accessible, that average; purple = near average accessibility or highly accessible). The most inaccessible gene is a HINT-proximal ferredoxin protein CCNA_03639. The most highly expressed mRNA in *Caulobacter, rsaA*, has average accessibility. These two genes are indicated in the plot. Some of the very most highly inaccessible genes do have higher than average expression levels, but in general there is no correlation (R^2^ = 0.070). The genes shown here exclude the two ribosomal RNA operons. RPKM is from (5), mixed population PYE.

**SI Figure 3.**
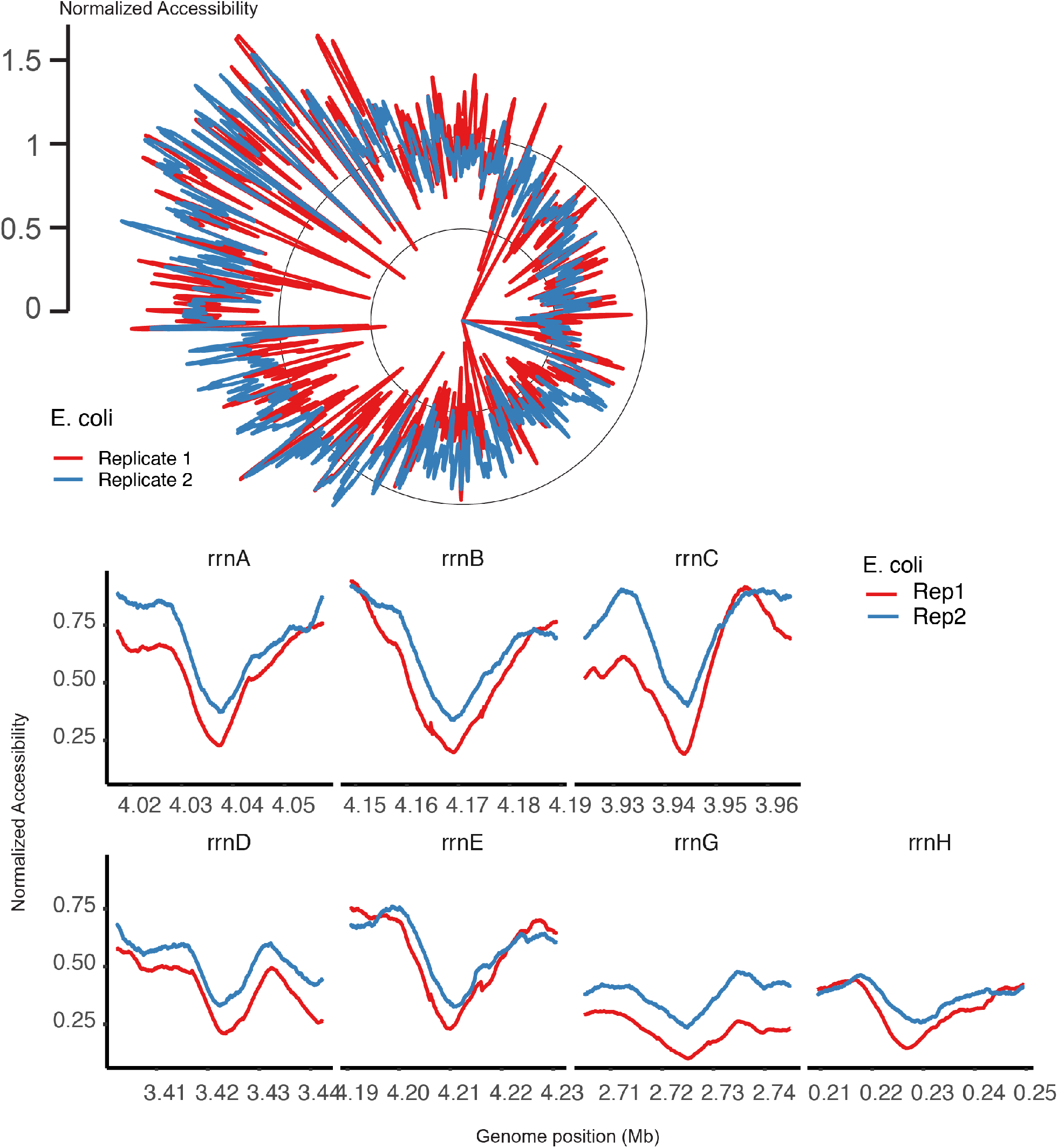
Bac-ATAC data for the *E. coli* chromosome in polar coordinates. For this Bac-ATAC data, coordinates are as described in Figure 1, but represent the *E. coli* chromosome rather than the *Caulobacter* chromosome. Data was obtained for plotting as described in SI Methods K. Upper Polar Plot: The origin is at ∼85 minutes and terminus is at ∼35 minutes, where the full chromosome (360 degrees) is 100 minutes. Locations where the ATAC-signal is lowest (closest to 0 radially) include every rRNA loci that was found to co-localize in *E. coli* cells in reference (18). This data was not normalized to genomic DNA and so is not corrected for copy number variation. Lower Cartesian Plot Panels: zoom-ins of the seven ribosomal RNA loci in *E. coli*. The y-axis represents normalized accessibility and the x-axis represents base pair coordinate in millions of base pairs. Red and blue lines represent data from replicate Bac-ATAC experiments. These regions are among the most highly inaccessible regions in *E. coli*, and therefore qualify as *E. coli* HINT regions.

**SI Figure 4.**
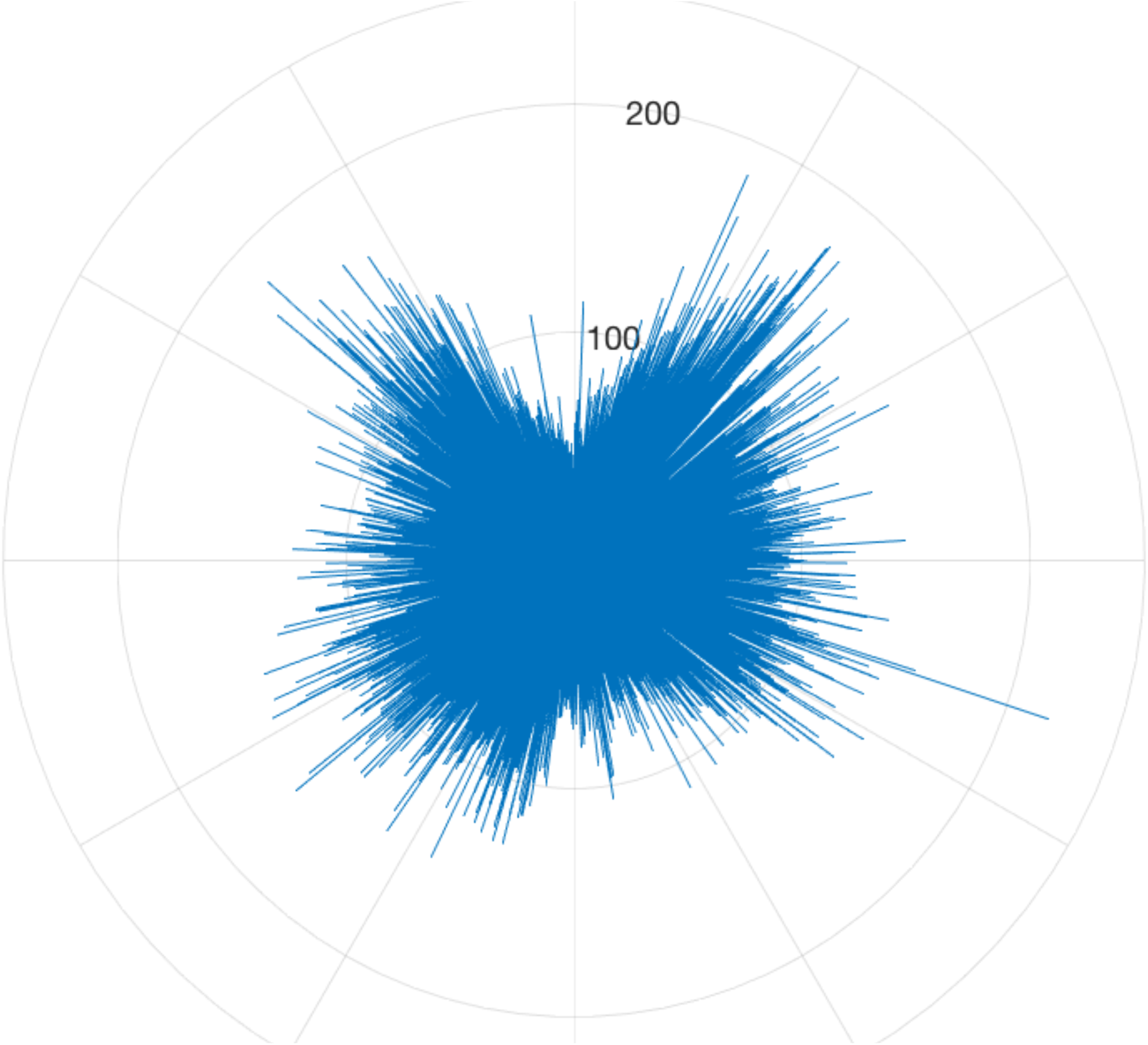
Raw insertion frequency data from Bac-ATAC on NA1000. Polar plot of raw insertion frequencies for WT data sample without normalization to genomic DNA or smoothening. Radial coordinate encodes frequency (each concentric grey circle represents an additional value of 100 mapped reads increasing radially) and the angular direction encodes base pair coordinate (with a resolution of 1bp).

**SI Figure 5.**
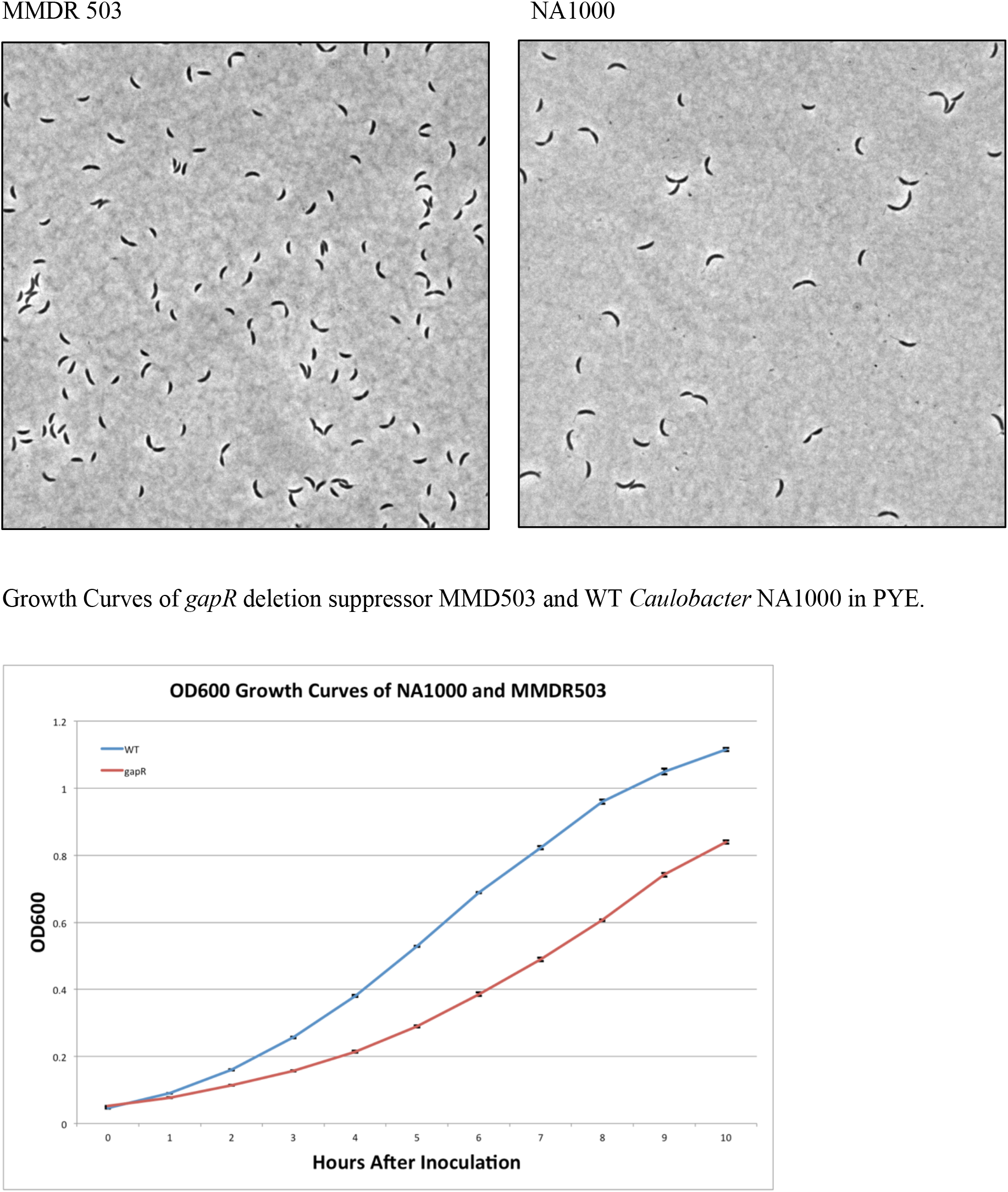
Micrographs of *gapR* deletion suppressors in PYE. Above: MMDR503 is a *gapR* deletion suppressor strain of NA1000 created using the protocol followed in **SI Methods 5**. Shown above are MMDR 503 (left) compared to wild type *Caulobacter* NA1000 (right). MMDR 503 was used for ATAC-seq, and was also whole-genome sequenced. MMDR 503 displays mostly WT cell morphology. Below: Comparative Growth Curves of *Caulobacter* NA1000 (blue) with MMDR503 *gapR* deletion suppressor in PYE.

**SI Figure 6.**
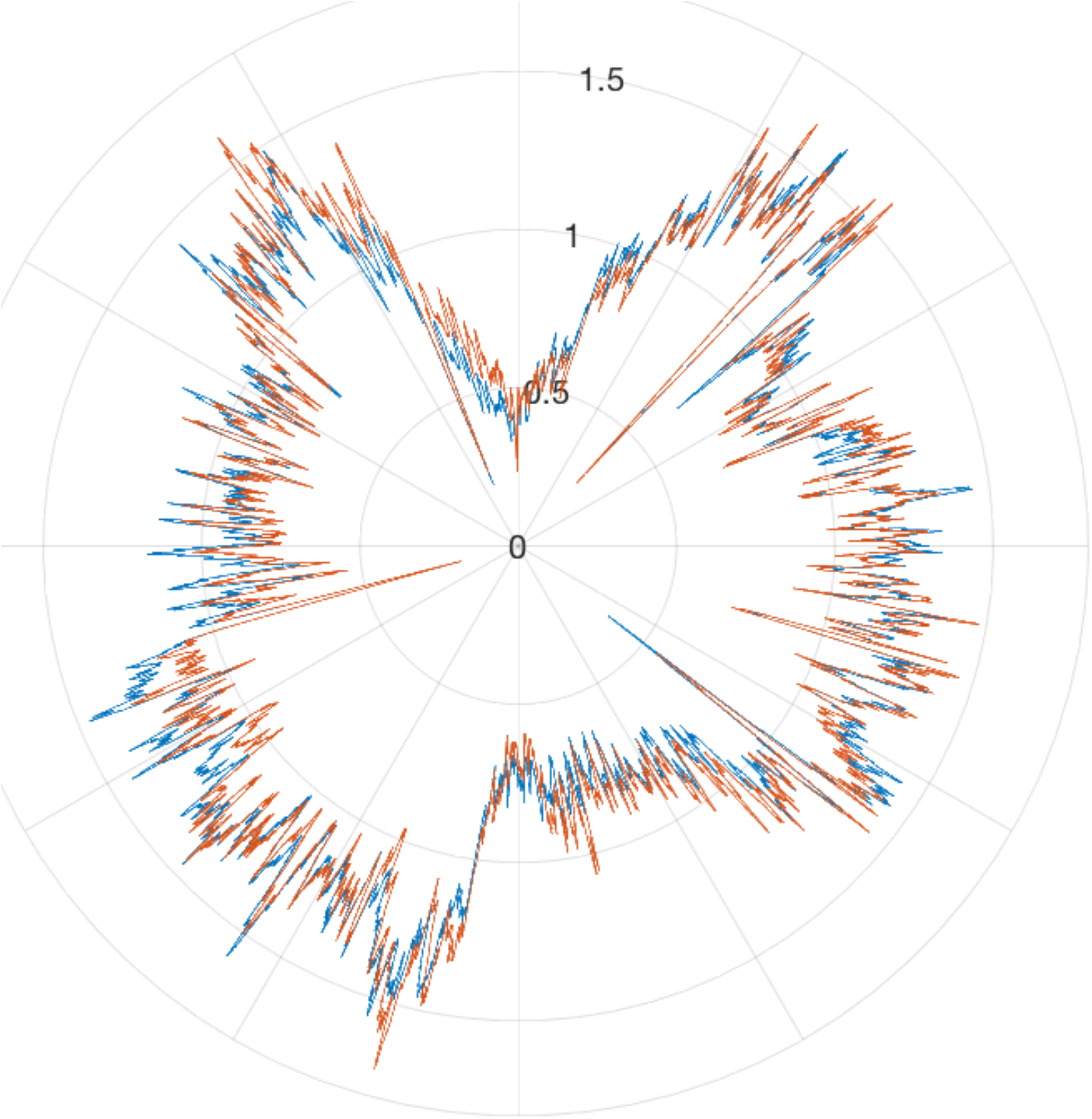
Bac-ATAC data from SMC deletion strain of *Caulobacter*. For this Bac-ATAC data, coordinates are as described in **Main Text Figure 4**, and data was obtained for plotting as described in **SI Methods K**, except there were no data points removed in this plot. Red line = SMC deletion mutant; Blue line = WT NA1000. WT genomic DNA data was used to control for copy number variations for both data sets. SMC is known to bind the origin in *Caulobacter*, where its ATPase activity is required for proper regulation of chromosome segregation (19). We performed ATAC-seq on SMC null mutant and observed that most of the global accessibility patterns of the chromosome are highly similar between NA1000 and the SMC deletion mutant.

**SI Figure 7.**
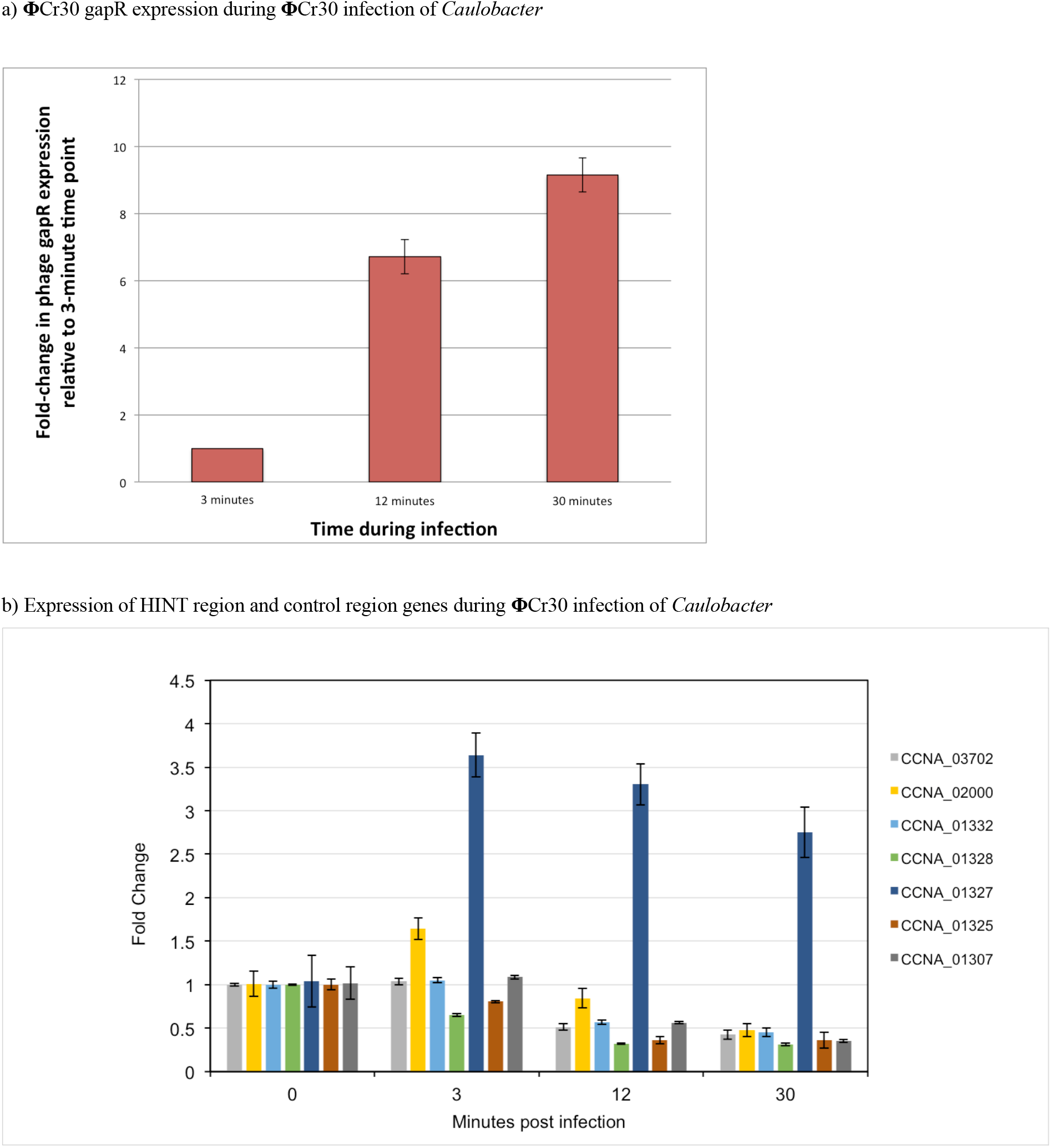
qPCR data during Phi ΦCr30 infection of *Caulobacter* mixed population. a) qPCR results showing that the **Φ**Cr30 ortholog of gapR is expressed during **Φ**Cr30 infection of *Caulobacter*. This transcript was not detected at t = 0 (immediately after infection initiation) so fold-changes are expressed in relation to expression level after 3 minutes of infection. Rho was used as an internal standard. b) Inaccessible genes in HINT 2 were tested for changes in transcription extent during phage infection. Bars show expression level fold-changes after specified lengths of time during infection. CCNA_03702 and CCNA_02000 serve as negative controls; these genes are ribosomal proteins located distal from the ribosomal protein operon HINT. CCNA_01327 is another control transcript, since this gene is actually found in the accessible region in the middle of HINT 2 (depicted as the peak at the center of HINT 2 in **Main Text Figure 1g**). The remaining genes are the experimental transcripts. CCNA_01307 is near the left edge of the HINT, whereas CCNA_01325 is at the rightmost edge of the first inaccessible operon associated with this HINT. CCNA_01328 and CCNA_01332 mark the left and right edges of the second set of inaccessible genes in this HINT (see **Main Text Figure 1g**). Whereas the inaccessible transcripts of this HINT region change similarly to the control genes CCNA_03702 and CCNA_02000 during infection, we note that the accessible transcript CCNA_01327 actually displays an increase in expression during phage infection.

**SI Figure 8.**
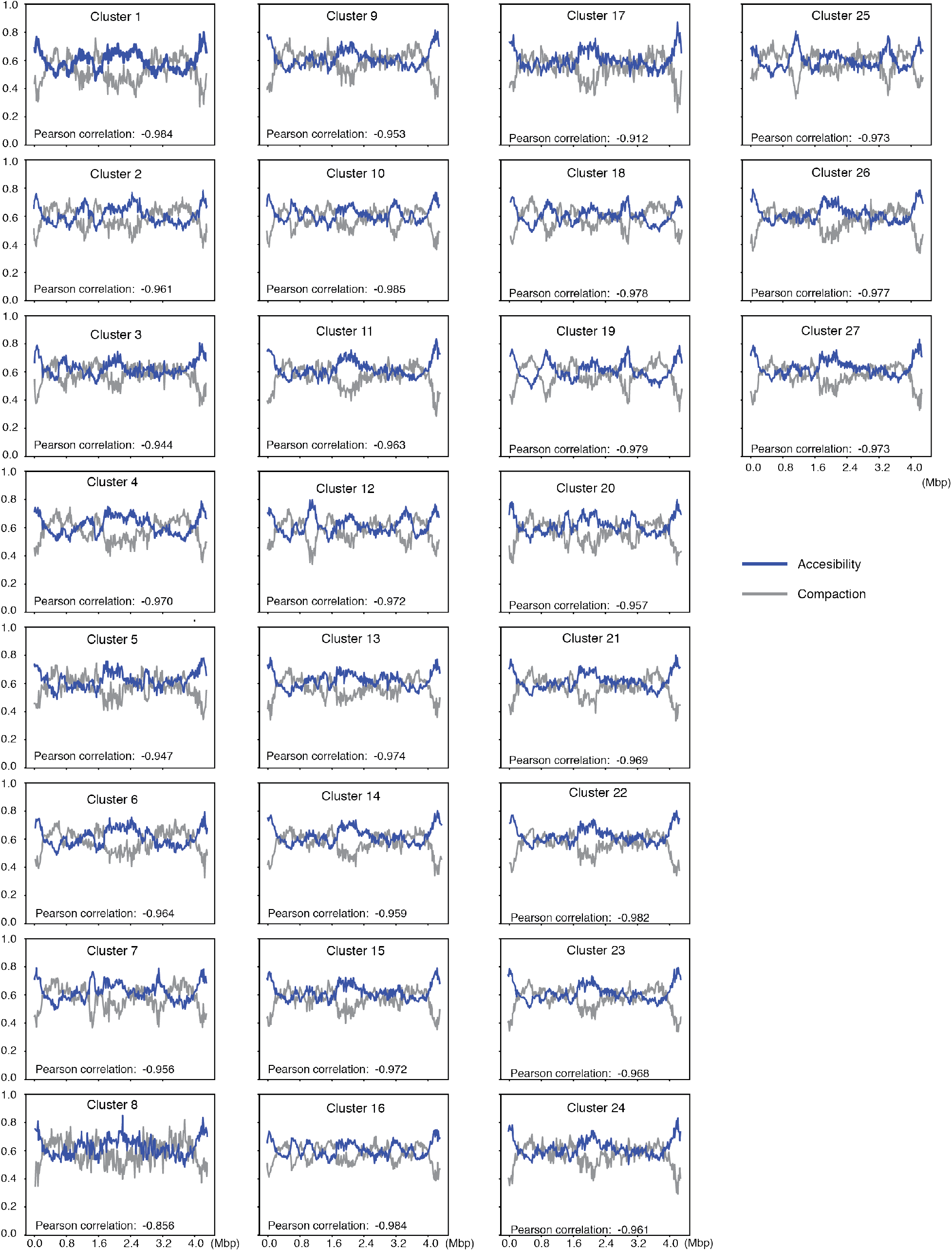
Anticorrelation of Accessibility and Compaction of DNA in computational models of the *Caulobacter* chromosome. The averaged accessibility (blue) and compaction (gray) scores are shown as a function of chromosomal position in 15000 bp bins. Pearson correlation is indicated for each cluster.

**SI Figure 9.**
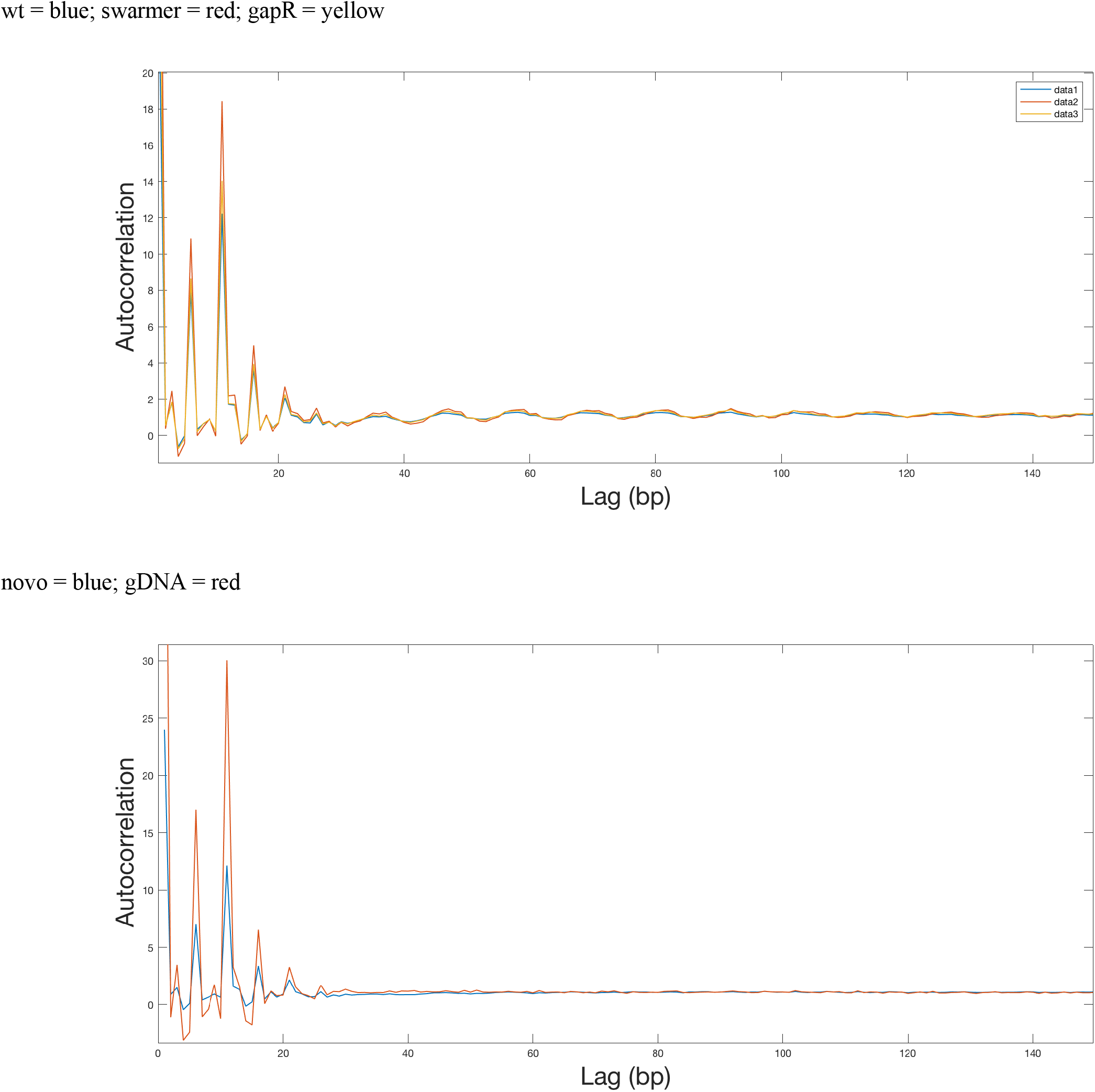
Autocorrelation Functions of Bac-ATAC data. Above: Autocorrelation function for Bac-ATAC data throughout the chromosome from zero to 150 bp of lag distance for the *Caulobacter* mixed population (blue), swarmer (red), and *gapR* deletion strain MMDR503 (yellow). Below: Autocorrelation function for Bac-ATAC data throughout the chromosome from zero to 150 bp of lag distance for novobiocin-treated *Caulobacter* (blue) and for purified genomic DNA (red).

**SI Figure 10.**
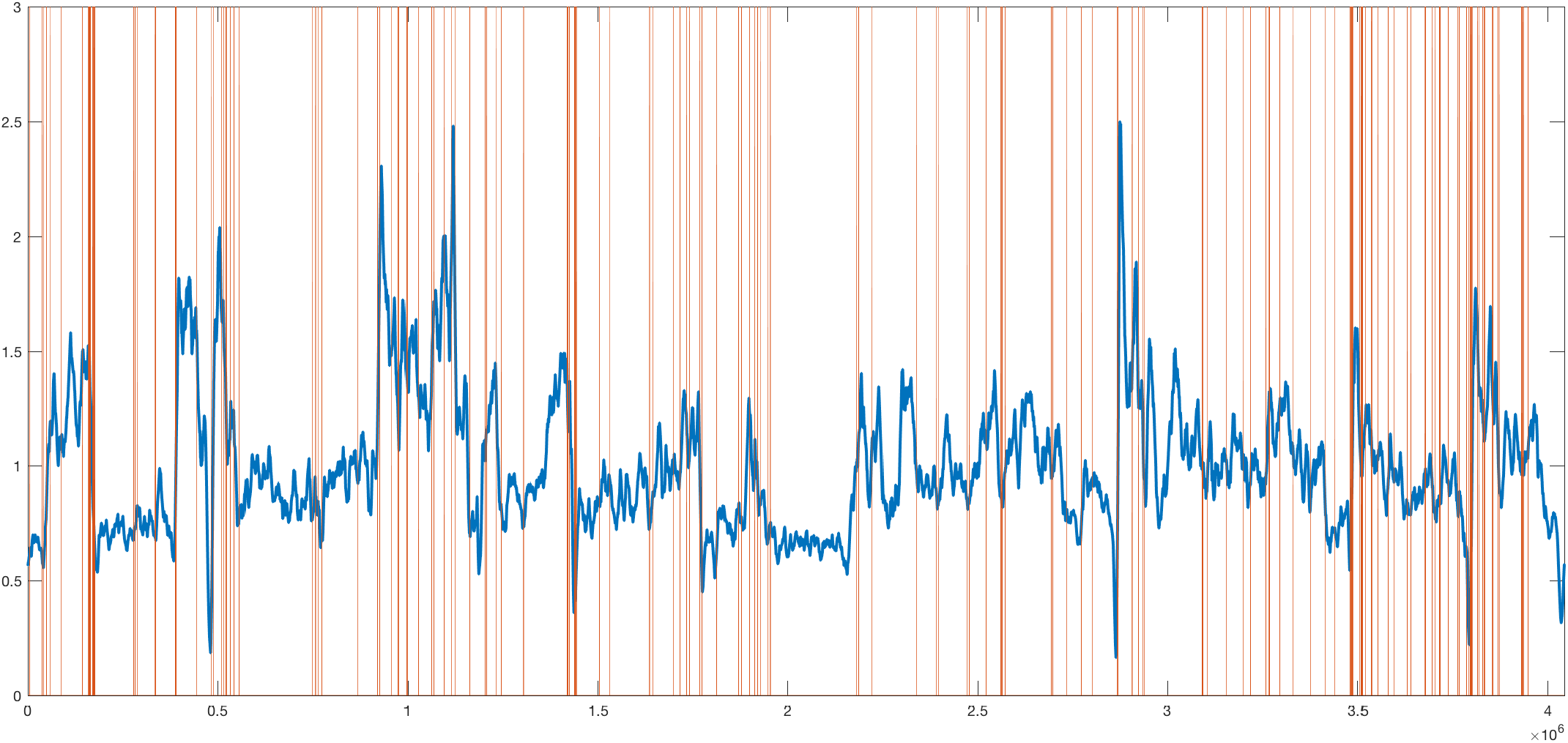
Novobiocin Bac-ATAC data overlaid with highly expressed genes in novobiocin. Y axis = Bac-ATAC normalized accessibility, smoothened over 10kb (blue) and x-axis is base pair coordinate of the chromosome (same depiction of data as in **Main Text Figure 1b**). Red lines represent the locations of the two hundred most highly expressed transcripts in *Caulobacter* (excluding antisense transcripts).

**SI Figure 11.**
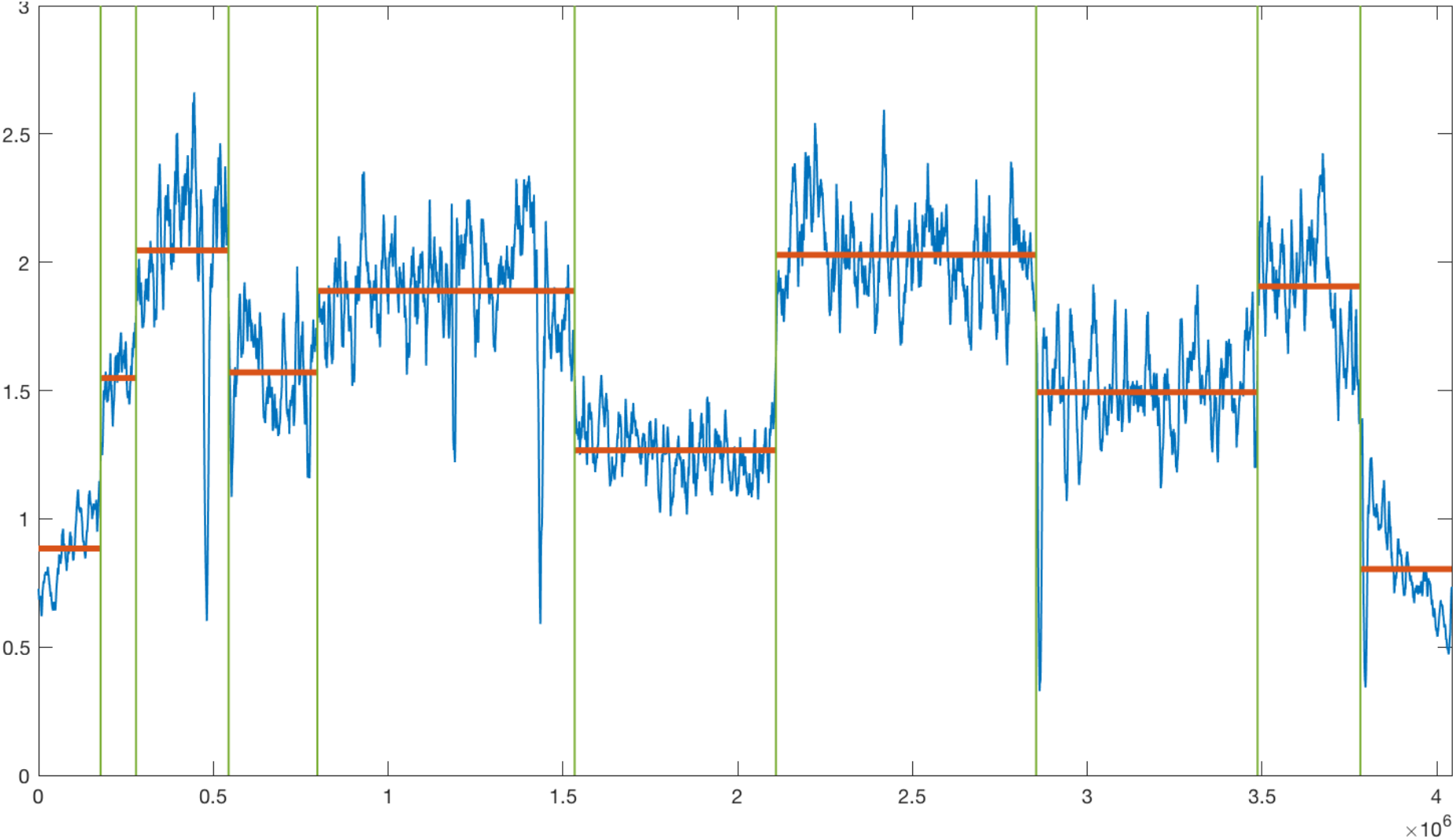
Initial results of change-point analysis on mixed population Bac-ATAC data. X-axis is bp coordinate, Y-axis is Bac-ATAC accessibility as in **Main Text Figure 1b** (blue line). This figure shows the raw output of **SI Methods 2**, where nine change-points were identified (green lines). The red lines show the running average accessibility within each resultant CAD. The second CAD boundary from the left was removed during analysis due to its proximity to its adjacent boundaries, for simplicity. The final list and description of the change-points (WT CAD boundaries) resulting from this analysis is in **SI Table 2**.

## Notes

https://www.ncbi.nlm.nih.gov/sra/PRJNA504229

### Competing Interest Statement

The authors have declared no competing interest.

